# Comparing trait syndromes between Taiwanese subtropical terrestrial and epiphytic ferns at the species and community level

**DOI:** 10.1101/2021.09.06.459074

**Authors:** Kenny Helsen, Jéssica Lira Viana, Tsung-Yi Lin, Li-Yaung Kuo, David Zelený

## Abstract

**Background and Aims:** While functional trait-trait and trait-environment relationships are well studied in angiosperms, it is less clear if similar relationships, such as the leaf economics spectrum (LES), hold for ferns, and whether they differ between terrestrial and epiphytic fern communities. We used vegetation data collected along an elevation gradient in Taiwan to explore these relationships.

**Methods:** We measured nine leaf traits for 47 terrestrial and 34 epiphytic fern species across 59 vegetation plots along an elevation gradient in the subtropical forest of Northern Taiwan. We explored trait-trait and trait-environment relationships at both the species and community levels for both growth habits, while accounting for phylogenetic relationships.

**Key Results:** Epiphytes differed from terrestrial ferns in species- and community-level trait values, mainly reflecting responses to higher drought and nutrient stress. The angiosperm LES was reflected in the trait-trait correlations of terrestrial ferns and less expressively in epiphytes. This pattern suggests that mainly water rather than nutrient availability shapes epiphytic trait patterns. Trait-trait analysis on raw trait data and on independent contrasts vary in some ways. Trait-environment relationships were similar for several drought-related traits across both species’ groups.

**Conclusions:** This study illustrates that fern trait patterns are not entirely equivalent for epiphytic and terrestrial species or communities and should not be extrapolated across growth habits or between the species and community levels. Phylogenetic constraints may influence the trait-environment response of epiphytic species.

## INTRODUCTION

Functional traits, *i.e.* morphological, physiological or phenological features that indirectly impact an organism’s fitness (Violle *et al*. 2007), often dictate under what environmental conditions plants will be able to thrive. This environmental trait filtering has been extensively studied and has increased our mechanistic understanding of species assembly at the community level (Lavorel and Garnier 2002; Kraft *et al*. 2015; Bruelheide *et al*. 2018; Nascimento da Costa *et al*. 2023). Several functional traits furthermore covary systematically across plant species. For example, the leaf economics spectrum (LES) links several leaf traits that govern a species’ leaf investment strategy. The spectrum ranges from highly resource-acquisitive, productive species characterised by high specific leaf area (SLA), leaf nitrogen and phosphorous content to more resource-conservative, slow-growing species with high leaf dry matter content (LDMC) and leaf lifespan (Wilson *et al*. 1999; Wright *et al*. 2004; Díaz *et al*. 2016). The LES is even retained when scaling up species-level traits to the community level (Bruelheide *et al*. 2018). The discovery of the LES and the links between LES traits and plant responses to several environmental stressors, such as low nutrient availability (Ordoñez *et al*. 2009; Hodgson *et al*. 2011) and low temperatures (Wright *et al*. 2005; Dong *et al*. 2020), have significantly increased our understanding of plant functioning and plant-environment interactions.

Most of this trait work has been performed on seed plants (spermatophytes), and it is unclear if the same trait-trait and trait-environment relationships hold for other vascular plant lineages, such as ferns (Karst and Lechowicz 2007). Since ferns are a distinct evolutionary lineage from seed plants (Smith *et al*. 2006), morphological and physiological traits are not guaranteed to be functionally equivalent across these groups (Peppe *et al*. 2014). For example, recent work has suggested that the evolution of photosynthesis-related trait co-regulations and trade-offs, such as the LES, has been more strongly constrained by leaf water transport capacity in ferns (Sessa and Givnish 2014; Zhang *et al*. 2014). The reason behind this is probably more limited potential for controlling evaporation, low water-use efficiency and xylem hydraulic limitations in ferns (Brodribb and Holbrook 2004; Zhang *et al*. 2014). Because of this, ferns are more vulnerable to drought stress, which may account for their relatively small range, at the more resource-conservative part of the global LES trait space (Díaz *et al*. 2016). Indeed, compared to angiosperms leaves, ferns have, on average, lower photosynthetic capability and leaf nutrient concentrations (Tosens *et al*. 2016). A few studies have nevertheless found some support for LES trait relationships across a limited number (*i.e.*, 11-35 species) of terrestrial ferns of temperate (Karst and Lechowicz 2007; Sessa and Givnish 2014; Tosens *et al*. 2016) and (sub)tropical ecosystems (Watkins *et al*. 2007; Zhu *et al*. 2016; Campany *et al*. 2019; Lin *et al*. 2020; Li *et al*. 2022), although with relative narrow scopes. Substantial evidence for global LES trade-offs among fern species, comparable to those reported for seed plants, is currently absent, as is an analysis contrasting LES patterns between growth habits. Our limited understanding of trait-trait relationships in ferns (cf. Kessler *et al*. 2016) is surprising, considering that they contain more than 11,000 species worldwide as the second largest lineage among land plants (Smith *et al*. 2006; PPG I 2016).

Almost one-third (29%) of all fern species are epiphytic (Schuettpelz and Pryer 2009; Dubuisson *et al*. 2009), with 61% of all epiphytic ferns from the Polypodiaceae and Hymenophyllaceae families (EpiList 1.0, Zotz *et al*. 2021). Especially in subtropical and tropical regions, ferns form the predominant vascular epiphytic group, together with orchids (Schellenberger Costa *et al*. 2018). The constraints of photosynthesis-related traits by water relationships are expected to be even stronger in epiphytic species since they are likely to experience more frequent and extended drought stress than terrestrial species (Dubuisson *et al*. 2009; Watkins and Cardelús 2012; Zhang *et al*. 2014; Aros-Mualin *et al*. 2021). Furthermore, epiphytes are confronted with low nutrient availability since they have no access to the soil (Watkins *et al*. 2007). On average, epiphytic ferns indeed have higher water-use efficiency (δ^13^C isotope ratio) and lower SLA, stomatal density and leaf N than terrestrial species in tropical forests, suggesting the importance of water and nutrient stress in shaping their traits (Watkins *et al*. 2007; Nitta *et al*. 2020; Campany *et al*. 2021). Consequently, it is unclear if similar LES-related trait relationships can be expected for epiphytic and terrestrial ferns (Hietz *et al*. 2022). For instance, a recent study on epiphytic plant communities along an elevation gradient in Tanzania found markedly different trait-environment correlations for epiphytic ferns and epiphytic flowering plants (Schellenberger Costa *et al*. 2018). Here, we employ phylogenetic data on trait syndrome studies since it is critical to understand the relationships among species and how character states differ between groups and clades (Ackerly 2009).

In ecosystems where terrestrial and epiphytic ferns co-occur, such as subtropical and tropical rain- and cloud forests, the environmental filtering of their respective communities might be governed by very different drivers. For example, the proneness to drought of epiphytes suggests that their community composition will be more strongly filtered by water availability through precipitation, relative air humidity, and ground fog cover (Dubuisson *et al*. 2009; Zhang *et al*. 2014; Aros-Mualin *et al*. 2021). Additionally, tree and shrub species composition directly affects the quality and availability of epiphyte substrate, thus further impacting their community composition. Terrestrial species, on the other hand, will also be filtered by soil variation through differences in soil nutrient and water availability, while the impact of temperature might be equally important for both growth habits. Several studies found that some morphological leaf traits vary in response to temperature (elevation) and humidity gradients for tropical terrestrial and epiphytic fern communities (Kessler *et al*. 2007; Kluge and Kessler 2007; Salazar *et al*. 2012; Petter *et al*. 2016). Nevertheless, studies exploring the importance of different environmental drivers on community-level trait composition of terrestrial and epiphytic ferns simultaneously for a given ecosystem are mainly lacking (however, see Wegner *et al*. 2003).

In this study, we measured nine functional leaf traits, including four LES-related traits, known to respond to nutrient, water and cold stress in seed plants, for a wide range of fern lineages, including 47 terrestrial and 34 epiphytic ferns across 59 plots. These plots were spread along an elevation gradient from 870 to 2130 m a.s.l. in the subtropical forests of Northern Taiwan. In addition to the measured species-level traits, we also calculated community mean trait values for each plot. Using this dataset, we explored the following research questions: 1) Are traits of terrestrial and epiphytic species and communities systematically different? 2) Do similar trait-trait correlations occur for terrestrial and epiphytic species and communities, and do these relationships mirror known trait patterns of seed plants, such as the LES? Are these correlations affected by phylogenetic relationships between species? 3) Do we find similar trait-environment relationships for terrestrial and epiphytic ferns?

## MATERIALS AND METHODS

### Study design

The study was conducted in the Wulai district of New Taipei City in northern Taiwan. In this area, we established 59 10 m × 10 m vegetation plots (Supplementary data Fig. S1) in natural, undisturbed vegetation along an elevational gradient ranging from Mount Meilu (870 m a.s.l., 24.85°N 121.53°E) to Mount Taman (2130 m a.s.l., 24.71°N 121.45°E). Vegetation along this gradient ranged from lowland *Pyrenaria-Machilus* subtropical winter monsoon forest to lower cloud zone *Quercus* montane evergreen broad-leaved cloud forest and *Chamaecyparis* montane mixed cloud forest (Li *et al*. 2013). The vegetation plots were evenly spread across a ridge in six elevation zones (with centres at 850, 1100, 1350, 1600, 1850 and 2100 m a.s.l. ±50 m), with ten plots per elevation zone spaced at least 50 m apart (except for the 1850 m zone, where only nine plots were located due to logistic constraints). At each elevation zone, plots were positioned along a topographic gradient across the ridge, ranging from the southwest-facing (leeward) to the northeast-facing slope (windward), including plots located on the ridge.

The climate in the study region is classified as a humid subtropical climate, with monthly mean temperatures in January and July ranging from 12.8 to 25.6 (near Mt. Meilu) and 8.7 to 21.1 (near Mt. Taman), respectively (obtained from on-site microclimatic measurements in a subset of our vegetation plots). Mean annual precipitation ranges from 2033 to 3396 mm along the gradient, with no obvious dry season (Supplementary data Fig. S2) (data from Lalashan and Fushan weather stations, respectively; Taiwan Central Weather Bureau, https://www.cwb.gov.tw). Microclimatic loggers installed at three sites at each elevation zone indicated high relative humidity for most plots (median between 97.5 and 100%). Parent materials of the study area mainly consist of Miocene and Oligocene sandstone and slate (Central Geological Survey, MOEA). The soils covering the parent material have high acidity and organic matter content, especially in higher elevations and on the ridges. Soils near ridges have higher degrees of podzolisation than those on steeper slopes due to lower degrees of soil layer disturbance on the flat microtopography of the ridges (Lin *et al*. 1988).

### Species composition and functional traits

In each plot, the understory terrestrial and epiphytic fern species composition was recorded (presence-absence data) from May to October 2017 and 2018, resulting in a species list containing 109 ferns. Nomenclature for species follows TPG (2019, 2021) (Supplementary data Table S1). Epiphytic species within reach were collected directly, while those out of reach were identified using binoculars and sampled for trait measurements using an 11.7 m long telescopic knife. We measured nine functional leaf traits for 47 terrestrial and 34 epiphytic ferns (63% and 79% of all recorded terrestrial and epiphytic ferns, including all large and abundant species, Supplementary data Table S1). For each of these species, we collected leaves from, on average, five (range 1-12) individuals directly in our plots or within a 50 m radius. Only adult individuals were selected, preferentially those capable of producing sporangia. For species occurring across different elevation zones, we attempted to collect individuals across this entire range. From each collected individual, we selected 1-6 leaves (fronds) for trait measurements, using the following criteria: 1) for dimorphic species, we selected the sterile rather than the fertile leaves; 2) the leaf should be fully expanded without visible herbivore or parasite-induced damage. For non-dimorphic species, traits were measured on sori-bearing fronds. Collected fronds were stored in wet, sealed plastic bags at low temperatures (< 10°C) for a minimum of 12h before trait measurement to allow full rehydration (cf. Pérez-Harguindeguy *et al*. 2013).

The measured traits consisted of leaf dry matter content (LDMC, mg g^-1^), specific leaf area (SLA, mm^2^ mg^-1^), leaf nitrogen content (leaf N, mg g^-1^), area-based leaf chlorophyll content (Chl, SPAD units), leaf area (LA, cm^2^), leaf thickness (Lth, mm), equivalent water thickness (EWT, mg mm^-2^), leaf ^13^C/^12^C stable isotope ratio (δ^13^C, ‰) and leaf ^15^N/^14^N stable isotope ratio (δ^15^N, ‰). EWT is sometimes known as ‘succulence’ (Mantovani 1999; Féret *et al*. 2019). The first four of these traits are part of the LES for vascular plants (cf. Wilson *et al*. 1999; Wright *et al*. 2004) and thus position species along a gradient from resource acquisitive (high SLA and leaf N) to resource conservative (high LDMC and area-based leaf Chl). The following four traits are expected to relate to drought stress, with small LA and high Lth, EWT and δ^13^C indicative of drought tolerance, with the latter trait a proxy of long-term water use efficiency (Farquhar *et al*. 1982; Medeiros *et al*. 2019; Maréchaux *et al*. 2020). δ^15^N reflects the nitrogen source used by a plant, with values around 0 ‰ usually indicating nitrogen fixation, while values around -2 and -6 ‰ indicating plant nitrogen acquisition through arbuscular and ectomycorrhiza, respectively (Craine *et al*. 2015). Trait measurements largely followed standard protocols (Pérez-Harguindeguy *et al*. 2013) but were partly adapted for fern leaves (Supplementary data Appendix S1, Fig. S3 - Fig. S6).

For all traits, except LA, values with a Z-score (standard score) larger than 2.5 at the species level were considered outliers caused by measurement error and were removed from the final dataset (<1% of data points). Next, all leaf-level traits were averaged to the species level, resulting in a 47 species × 9 traits matrix for the terrestrial species and a 34 species × 9 traits matrix for the epiphytic species. Missing leaf Chl values of four epiphytic species were replaced by the mean Chl of all species belonging to the same (sub)family. Trait values were also translated to the plot level by calculating community mean (CM) trait values, *i.e.,* the average trait value across all species in the plot, not weighted by species abundances, since these were not collected. CMs were first calculated from the original non-standardised traits (‘cwm’ function from *weimea* R package, Zelený 2020) and then standardised to Z-scores. Leaf area, Lth and EWT were logarithmically transformed in all species combined dataset before species- and community-level calculation to address the strong data skewness.

### Climate proxies

In each plot, we recorded the exact elevation using a topographic map combined with the GPS coordinates (GPSMAP 64st, Garmin, USA), aspect using a compass (SILVA, Sweden), and slope using a clinometer (SUUNTO PM-5/360 PC Clinometer, SUUNTO, Finland). We then calculated the heat load from the aspect folded on the prevailing wind direction (45°) and the slope with equation 2 of McCune and Keon (2002). Heat load quantifies the potential temperature increase that the plot can obtain through absorbing incident solar radiation.

Using the ground fog frequency raster map for Taiwan, we determined the average annual ground fog frequency for each plot (Schulz *et al*. 2017; 250 m spatial resolution). Variation inflation factors (VIF) indicated that the three climate proxies, namely elevation, fog frequency and heat load, were largely independent of each other (VIF < 1.5) (Zuur *et al*. 2010). VIFs were calculated with the ‘vif’ function in the *car* R package (Fox and Weisberg 2019).

### Species-level trait-trait relationship

Differences in average trait values between growth habits were assessed with a Mann-Whitney *U* test for each trait. Pairwise trait-trait relationships within growth habits were estimated with Spearman rank correlations. Additionally, we visualised the multivariate trait variation through principal component analyses (PCA) performed with the *vegan* R package (Oksanen *et al*. 2019). We applied three PCA analyses on all species combined, only terrestrial species, and only epiphytic species. Furthermore, we constructed separate trait hypervolume analysis for only terrestrial and only epiphytic species, using the first three axes of the PCA for all growth habits combined, using the *hypervolume* R package (Blonder *et al*. 2014; Blonder 2018) and following the protocol described in Helsen *et al*. (2020). These hypervolumes were used to quantify the trait space size and overlap between terrestrial and epiphytic species.

To account for the shared evolutionary history of species while performing species-level trait-trait relationship analyses (Felsenstein 1985), we repeated the pairwise correlations and PCA analysis within a comparative phylogenetic framework. First, we reconstructed a dated phylogeny based on the FTOL v1.1.0 dataset (Fern Tree of Life, Nitta et al. 2022), which we expanded with ten species to cover the full species list of our study (See Supplementary data Appendix S2 for details). The resulting dated tree (with 5568 species) was pruned to show relationships among the species in our study (81 species). We estimated phylogenetic independent contrasts (PIC) for each continuous trait with the “pic” function in the R package *ape* (v 5.0, Paradis and Schliep 2019). Employing raw trait or PIC data, pairwise correlations and PCA analyses were performed in the same manner. For pairwise correlations with PIC data, the exception was that rather than Spearman rank correlation, we used Pearson product moment forced through the origin. Phylogenetic stochastic mapping methods were used to reconstruct growth habit state changes across our phylogenetic tree. We performed maximum likelihood ancestral state reconstructions for growth habit using the “ace” function in the R package *ape*. Shifts among states across nodes were fitted with an equal-rates model. Nodes with scaled likelihood values for the terrestrial state higher than 0.5 were assigned to terrestrial ferns, otherwise epiphytes (Supplementary data Fig. S7). We acknowledge that phylogenetic uncertainty was not addressed when estimating state transitions with ancestral state reconstruction (Duchêne and Lanfear 2015).

We also estimated Pagel’s lambda (λ) (Pagel 1999) for each trait with function ‘phylosig’ of the *phytools* R package (Revell 2012). Pagel’s λ ranges between 0 and 1. When λ approaches one, it indicates a strong phylogenetic signal following a Brownian Motion model of evolution. Statistical significance of λ was determined by log-likelihood ratio tests comparing the predicted λ to a null model.

### Community-level trait-trait relationship

Community-level trait-trait relationship was analysed similarly to species-level trait-trait analysis, using CM trait matrices instead of species-level trait. We tested differences in average trait values between growth habits with Mann-Whitney *U* tests. Pairwise trait-trait relationships were tested by Spearman’s rank correlation. Since CM values are derived from species trait values and species composition data, CM values calculated for different traits but from the same species composition data (*e.g*., for epiphytes) are not independent, and their correlation cannot be tested by standard tests expecting independence of variables. To avoid the problem of spurious correlation (Pearson 1897; Zelený 2018), we applied a modified permutation test using species-level permutation (comparing test statistics calculated from observed CM values with the distribution of test statistics calculated from CM values derived from species trait values permuted 49,999 times between species). Additionally, for each trait, we also performed a Spearman’s rank correlation between epiphytic and terrestrial species CM traits. Multivariate trait variation was visualized though three PCA analyses on the standardized plot × CM trait matrices (one for all species combined, one for only terrestrial species and another for only epiphytic species). CM trait space size and overlap for both species’ groups was again assessed through trait hypervolume construction, based on the first three axes of the PCA performed on the full plot × CM trait matrix.

Significance levels of all analyses, including all trait-trait correlations and average trait comparisons between species groups at both the species and community levels, were adjusted for Type I error inflation due to multiple testing using the false discovery rate method (Benjamini and Hochberg 1995) with the ‘p.adjust’ function in the *stats* R package.

### Community-level trait-environment relationships

To explore relationships between the measured traits and the three climate proxies (elevation, fog frequency and heat load), we used the fourth-corner approach, which calculates (weighted) Pearson correlations between a single trait from the standardised species × trait matrix and a single environmental variable from the plot × environmental data, weighted by the plot × species matrix (Legendre *et al*. 1997; Dray and Legendre 2008; ter Braak *et al*. 2018). We computed the original and Chessel fourth-corner correlation coefficients, following the recommendations of Peres-Neto *et al*. (2017). The latter is the original correlation coefficient divided by its maximum attainable value, which measures how well the trait-environment correlation explains the species distribution (Peres-Neto *et al*. 2017). The fourth corner correlations were tested by the ‘max test’ proposed by ter Braak *et al*. (2012), which overcomes Type-I error inflation issues of trait-environment correlations (ter Braak *et al*. 2012, 2018), using the function ‘test_fourth’ in the *weimea* R package (Zelený, 2018; https://github.com/zdealveindy/weimea). All analyses were performed in R, version 4.2.2 (https://www.r-project.org/).

## RESULTS

### Species-level analysis

Comparing the traits of the 47 terrestrial and 34 epiphytic ferns in our study showed that, on average, epiphytic species had lower SLA, LA, δ^15^N, leaf N, and higher EWT, and δ^13^C than terrestrial species (Fig. 1, Supplementary data Table S2). These differences were reflected in the PCA trait space, where epiphytic and terrestrial species were largely segregated (Fig 2A, Supplementary data Table S3A), and thus both had a relatively large proportions of unique trait space (61.62% and 72.94% respectively), following the hypervolume construction (Supplementary data Table S4, Fig. S8). The epiphytic species trait hypervolume was furthermore 1.45 times larger than the terrestrial species trait hypervolume, despite containing less species. This seemed to be mainly caused by a larger variation in EWT, Lth, Chl, and LDMC values among epiphytic species, compared to terrestrial species (Fig. 1). Standard PCA and phylogeny corrected PCA showed similar trait relationships patterns. In both PCAs, EWT, Lth, Chl and SLA contributed with a large proportion of the variation along the first PC axis (Supplementary data Fig. S9A, B; Table S3B).

**Figure 1.**
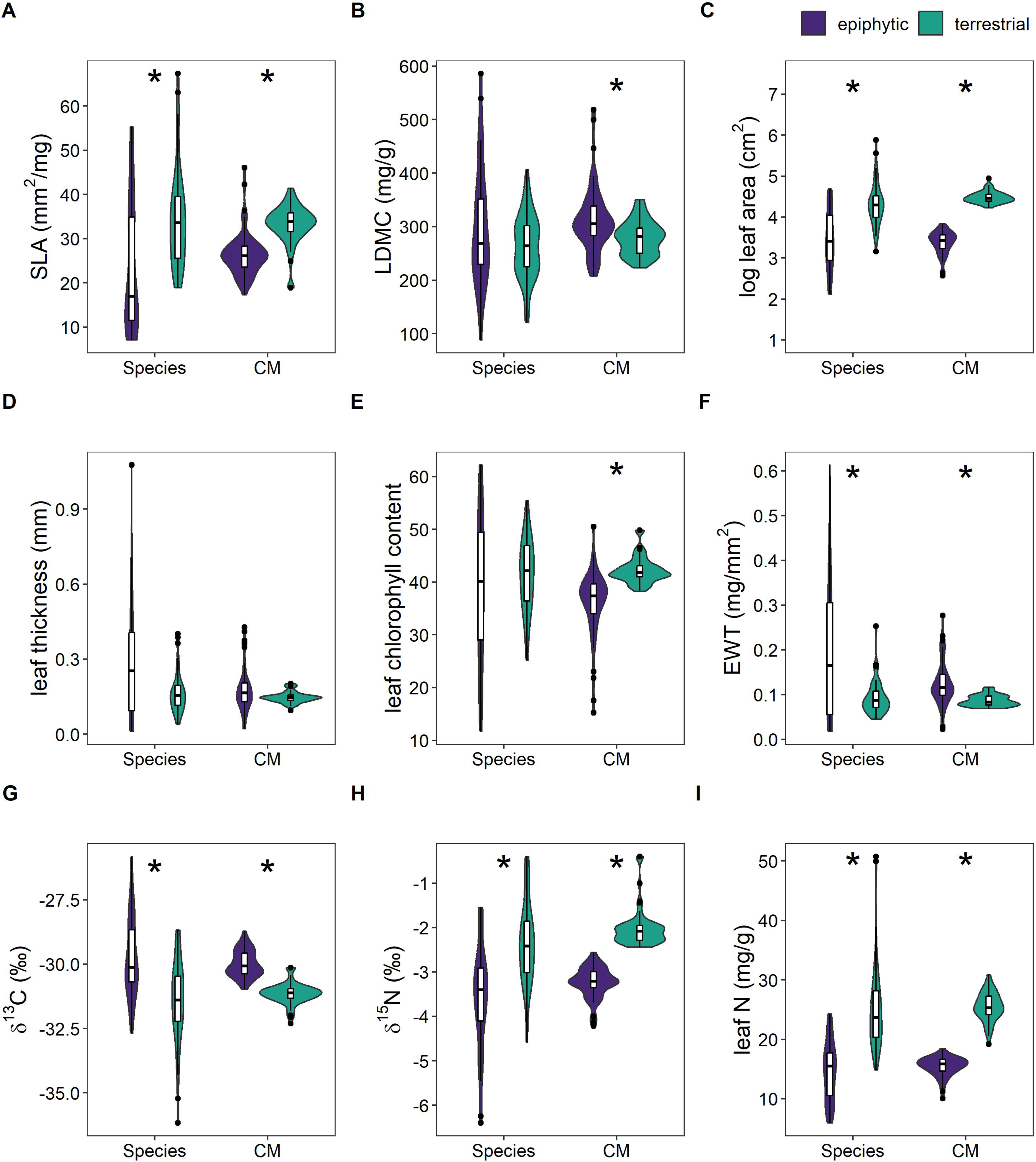
Violin plots for each trait separately, at both the species and community (CM) levels. Separate plot for epiphytic (dark purple) and terrestrial species (light green). Asterix indicates a significant difference between epiphytic and terrestrial growth habits at either species or plot levels (Supplementary data Table S2).

**Figure 2.**
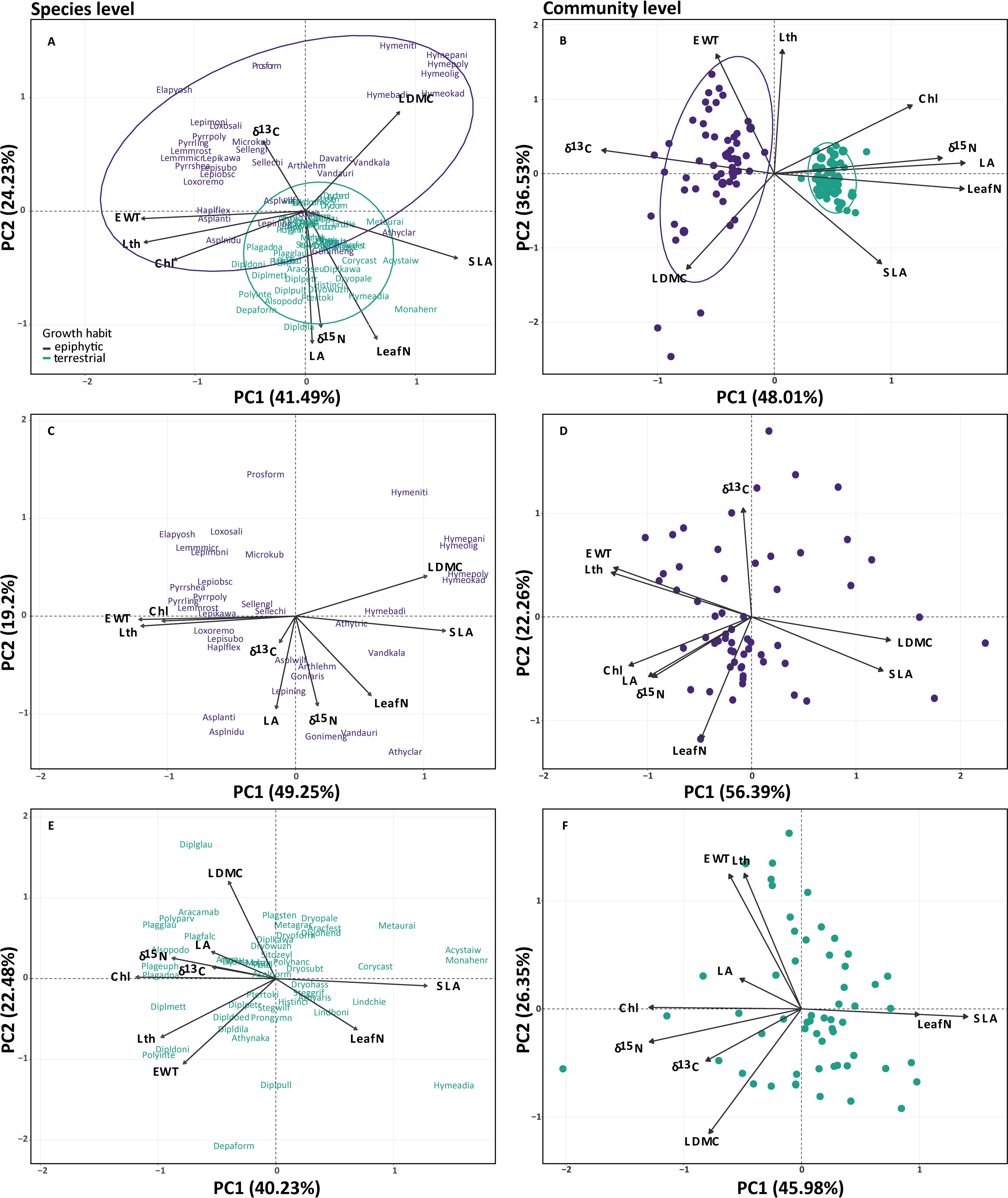
Biplots for the principal component analyses done on raw trait data (left column) and CM of traits (right column). A. the full species × trait matrix, B. plot × community mean (CM) trait matrix using all species combined, C. epiphytic species × trait matrix, D. plot × epiphytic CM trait matrix, E. terrestrial species × trait matrix, and F. plot × terrestrial CM trait matrix. Chl = leaf chlorophyll content, δ^13^C = the leaf ^13^C/^12^C stable isotope ratio, δ^15^N = the leaf ^15^N/^14^N stable isotope ratio, EWT = equivalent water thickness, Lth = leaf thickness, LA = leaf area, Leaf N = leaf nitrogen content, and LDMC = leaf dry matter content. For PCA on phylogenetic independent contrasts, see Fig. S9.

Different patterns emerged when comparing trait patterns separately for epiphytic and terrestrial species. LDMC was positively correlated with SLA and Chl suggesting a partial disconnection from the expected LES links (Supplementary information Fig. S10). However, the strength and direction of these relationships changed, and the expected LES trait correlations became more apparent in the phylogeny-informed analysis. Epiphytic ferns obtained more significant correlations than terrestrial ferns due to both SLA and LDMC links to drought-related traits. SLA and drought tolerance traits obtained stronger coefficients than with other leaf economics traits when using PICs (Supplementary data Fig. S11). In standard PCA, epiphytes seemed to cluster in three more-or-less distinct groups in the ordination space. The first group showed high-LDMC species, the second group showed high values for EWT and Lth values, and the third showed less extreme trait values overlapping the terrestrial species trait space (Fig. 2A). Despite the observed changes in the direction and strength of the relationships among SLA, LDMC and Chl, we found that trait patterns in epiphytes were generally similar between standard and phylogeny-informed analyses (Supplementary data Fig. S9 C,D).

Terrestrial species showed trait correlations expected under the LES links only in standard analysis not accounting for phylogeny. After phylogenetic correction, the LES trait links became less defined with no significant correlations between LDMC and SLA or Chl. Still, the negative correlation between leaf N and δ^13^C suggest that acquisitive terrestrial ferns with high values of leaf N were less water efficient (higher negative values of δ^13^C), the opposite pattern was observed in epiphytes. In the standard PCA, terrestrial species were separated by high SLA and high LDMC species occupying the opposite ends of the ordination trait space gradient. An additional group of species exhibited large EWT, Lth, and intermediate SLA and LDMC values (Fig. 2E). Phylogeny corrected PCA and Pearson correlations exhibited similar trait patterns, showing strong relationships among SLA EWT and Lth. In addition, relationships among Chl, δ^13^C and δ^15^N which had been significantly positive in standard analyses disappeared (Supplementary data Fig. S8 F, S10).

We detected a high degree of phylogenetic signal expression in the functional traits for these subtropical fern species. All leaf traits showed a significant phylogenetic signal, with fitted lambda values ranging from 0.55 to 0.97 (Supplementary data Table S5).

### Community-level analysis

With a few exceptions, community-level trait patterns were generally similar to those found at the species-level. For instance, compared to terrestrial species, epiphytes showed higher CM LDMC and lower CM Chl (Fig. 1B, E), while CM Lth did not differ (Fig. 1D). In parallel with species-level, proportions of unique trait space of epiphytes were greater than terrestrial ferns (93.15% and 88.48%, respectively), and the difference in hypervolume size between the species groups (1.87 times larger for epiphytes) were slightly more pronounced at the community-level (Supplementary data Table S4, Fig. S8). Note that the constructed three-dimensional hypervolumes are larger than the ellipses visualised in Fig. 2A, B due to their probabilistic nature. In general, the trait space of terrestrial and epiphytic ferns was completely partitioned at the community level (Fig. 2B). Comparison between standard PCA for species and communities showed similar trait patterns, with epiphytes showing consistently higher δ^13^C, lower SLA and leaf N values (Fig. 2B, Supplementary data Table S6).

Pairwise trait correlations patterns in the community level were similar to those found in standard species-level analysis for both epiphytes and terrestrials. However, community-level patterns did not completely mirror those at the species-level. For example, significantly positive correlation between Chl and LA for epiphytes, and Chl and δ^15^ N for terrestrials were only observed at the community level (Supplementary data Fig. S12). Significant trait correlations involving LMDC at the standard species level analysis were more strongly linked at the community level for epiphytic species, while for terrestrial species communities they completely disappeared. Community-level traits were furthermore only positively correlated (ρ**_s_**, *P* < 0.05) between epiphytic and terrestrial species across the vegetation plots for LDMC, EWT and Lth (Fig. 3, Supplementary data Table S7). At the same time, δ^13^C was negatively correlated between growth habits.

**Figure 3.**
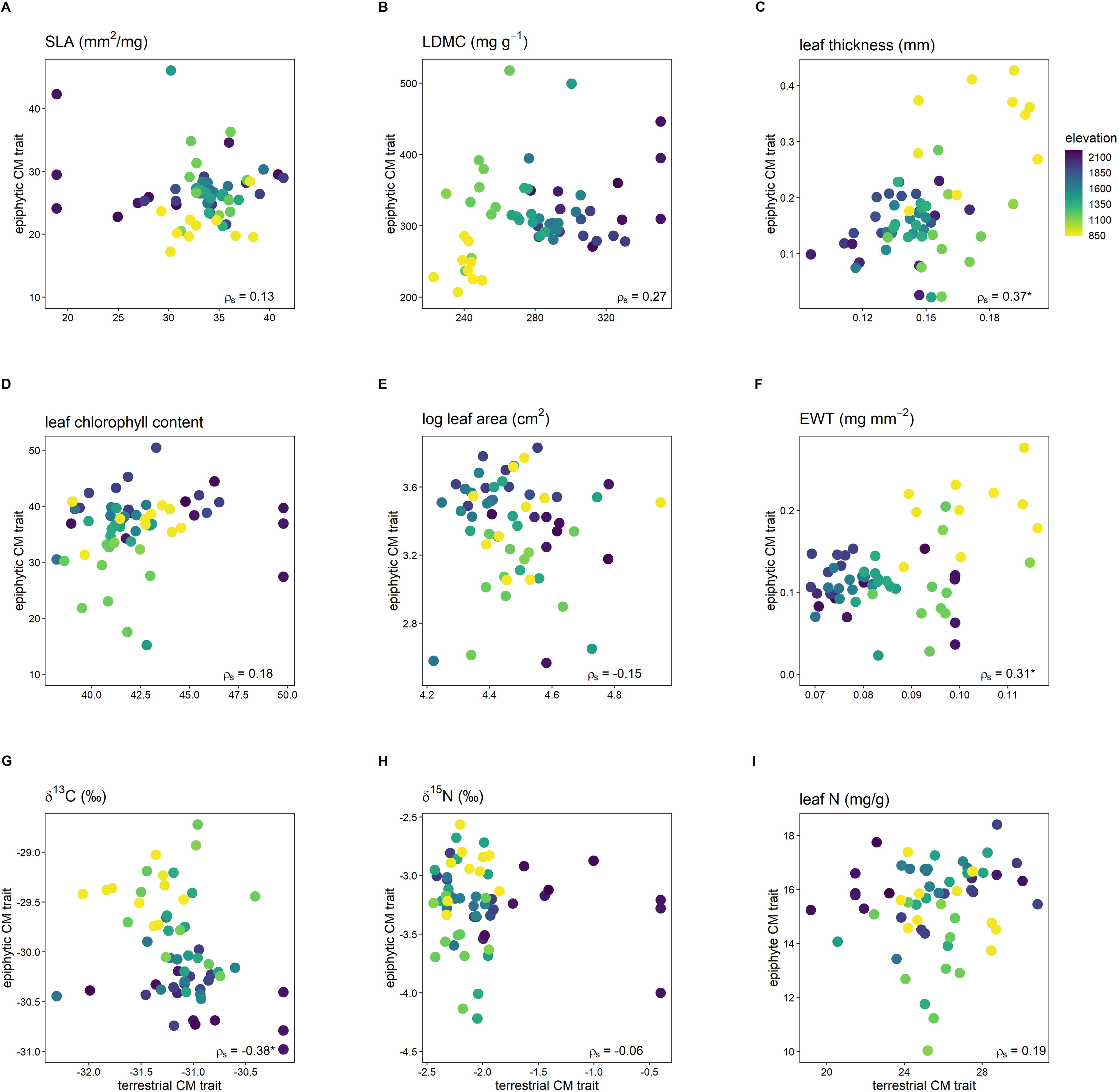
Scatterplots for pairwise Spearman’s rank correlation between plot-level terrestrial and epiphytic fern community mean (CM) trait values. Each data point corresponds to one vegetation plot, with colours indicating plot elevation. δ^13^C = the leaf ^13^C/^12^C stable isotope ratio, δ^15^N = the leaf ^15^N/^14^N stable isotope ratio, EWT = equivalent water thickness, Leaf N = leaf nitrogen content, and LDMC = leaf dry matter content. ρ_s_ = Spearman’s rank correlation coefficient. * = P-adj. < 0.05.

### Trait-environment relationships

The responses related to trait-environment relationships were not completely similar for both growth habits. While there were several significant (*P* < 0.05) relationships between traits of terrestrial species and environmental variables, traits of epiphytic species did not show significant responses. For terrestrial species, EWT was negatively related to elevation and fog while LDMC showed a positive response for these two climatic proxies. Leaf LDMC was also negatively related with heat load. Trends in trait-environment relationships in respect to epiphytic species was observed between δ^13^C and elevation (negative), and between δ^15^N and heat load (positive), however, these trends were not significant after FDR correction (Table 1).

**Table 1.** Fourth-corner results for individual trait–environment relationships for epiphytic and terrestrial species separately.

|  | elevation |  | fog |  | heat load<sup>sq</sup> |  |
| --- | --- | --- | --- | --- | --- | --- |
|  | epiphytic | terrestrial | epiphytic | terrestrial | epiphytic | terrestrial |
| SLA | 0.08/0.11 | 0.00/0.00 | 0.04/0.06 | 0.03/0.04 | -0.05/-0.07 | 0.03/0.05 |
| LDMC | 0.13/0.18 | 0.39/0.51<sup>*</sup> | 0.06/0.08 | 0.29/0.38<sup>*</sup> | -0.11/-0.15 | -0.12/-0.16<sup>*</sup> |
| Leaf area<sup>log</sup> | 0.08/0.10 | -0.03/-0.04 | 0.11/0.15 | -0.07/-0.10 | 0.01/0.02 | -0.02/-0.03 |
| Leaf thickness<sup>log</sup> | -0.13/-0.18 | -0.19/-0.25 | -0.07/-0.09 | -0.14/-0.18 | 0.10/0.13 | 0.05/0.06 |
| Chlorophyll content | 0.14/0.19 | 0.03/0.04 | 0.10/0.13 | -0.01/-0.01 | 0.00/0.01 | -0.04/-0.05 |
| EWT<sup>log</sup> | -0.13/-0.18 | -0.30/-0.39<sup>*</sup> | -0.07/-0.10 | -0.26/-0.35<sup>*</sup> | 0.09/0.12 | 0.05/0.07 |
| $\delta^{13}\text{C}$ | -0.24/-0.33 | 0.07/0.10 | -0.09/-0.12 | 0.05/0.07 | 0.04/0.05 | -0.08/-0.11 |
| $\delta^{15}\text{N}$ | -0.03/-0.04 | 0.07/0.10 | 0.02/0.04 | -0.03/-0.04 | 0.11/0.15 | -0.05/-0.07 |
| Leaf N | 0.08/0.11 | 0.02/0.03 | 0.11/0.15 | 0.02/0.03 | 0.01/0.02 | <0.01/0.01 |

Fourth-corner and Chessell-transformed correlation coefficients (r/Chessell-transformed r). $\delta^{13}\text{C}$ = the leaf $^{13}\text{C}/^{12}\text{C}$ stable isotope ratio, $\delta^{15}\text{N}$ = the leaf $^{15}\text{N}/^{14}\text{N}$ stable isotope ratio, EWT = equivalent water thickness, LDMC = leaf dry matter content, leaf N = leaf nitrogen content, SLA = specific leaf area. <sup>sq</sup> = squared transformation, <sup>log</sup> = logarithmic transformation. Codes for p-values after FDR correction, $< 0.05 = *$ .

## DISCUSSION

### Trait differences between epiphytic and terrestrial species

Epiphytic species were functionally differentiated from terrestrial species in our study, with most measured leaf traits significantly different, at both the species and community levels. These differences agree with fern studies in other (sub)tropical regions in Asia, Central America, and Polynesia. Higher Lth, EWT and δ^13^C in epiphytic species are usually attributed to the higher frequency and intensity of drought events experienced by these species, compared to terrestrial ferns (Watkins *et al*. 2007; Campany *et al*. 2021; Hietz *et al*. 2022). In addition, small size (*i.e.,* lower LA) for epiphytes has also been linked to water relations in a study in French Polynesia (Nitta *et al*. 2020; Jin *et al*. 2021), although several other studies found no significant difference in LA between epiphytic and terrestrial ferns (Zhang *et al*. 2014; Campany *et al*. 2021). The importance of drought stress for epiphytic ferns is further supported by their lower stomatal density observed in other studies (Zhang *et al*. 2014; Campany *et al*. 2021).

Other trait differences, such as low SLA and leaf N in epiphytes compared to terrestrial species, are also consistent with previous studies (Watkins *et al*. 2007; Zhang *et al*. 2014; Nitta *et al*. 2020; Campany *et al*. 2021) and reflect a shift along the LES towards more resource conservative strategies for epiphytes (Wright and Cannon 2001; Wright *et al*. 2004). This shift is likely driven by the lower nutrient availability experienced by epiphytes compared to terrestrial species (Watkins *et al*. 2007). However, SLA differences could also be indirectly caused by water stress avoidance (Campany *et al*. 2021). The lower δ^15^N signature of epiphytes has been hypothesised to reflect the uptake of ^15^N-depleted, atmospherically derived N sources through precipitation and fog (Watkins *et al*. 2007; Craine *et al*. 2015). Higher LDMC and lower leaf chlorophyll at the community level for epiphytes might also reflect more resource-conservative strategies and suggest that epiphytes with this set of traits are more common at the community level (Wright *et al*. 2004; Hodgson *et al*. 2011).

Trait variation was considerably larger among epiphytes than among terrestrial ferns at both species and community levels, reflecting patterns from previous studies (Watkins *et al*. 2007; Nitta *et al*. 2020). This large variance found in epiphytes seems to reflect the co-occurrence of three more-or-less distinct drought-coping trait syndromes (Kessler *et al*. 2007), with species belonging to phylogenetic distinct groups. The first group consists of species with very high LDMC and very low Lth, which are commonly associated with the filmy ferns (family Hymenophyllaceae). Species with high values of LDCM such as *Hymenophyllum badium* and *H. polyanthos* can completely dry out when under water stress and rehydrate after a drought period passes (Garcés Cea *et al*. 2014). Kessler *et al*. (2007) classified this as a ‘poikilohydric’ drought strategy. The second group, termed ‘xeromorphic’ by Kessler *et al*. (2007), contains species that prevent desiccation through thick, fleshy and small leaves with high Lth and EWT, such as *Lemmaphyllum microphyllum*. The bird’s nest ferns *Asplenium antiquum* showed a similar strategy, although bearing larger leaves. The third group has more intermediate trait values, with seemingly no pronounced drought-related traits. This group seems to merge species with two main drought-related strategies. Firstly, mesomorphic species that avoid drought by growing in the moistest microclimates, such as *Lepidomicrosorium ningpoense*, and secondly, species that have succulent rhizomes and shed their leaves to reduce water stress during drought (drought-deciduous species). The latter strategy is observed both in *Goniophlebium amoenum* and *Selliguea echinospora*. Higher trait variation among epiphytic species has been attributed to the strong vertical gradients in light intensity, temperature and humidity occurring in forest (Petter *et al*. 2016). This allows epiphytes to sort vertically in different niches from the dark and humid understory to the sunny and dry outer canopy, thus presenting a more varied environment than experienced by terrestrial species, which can only sort horizontally along the forest floor (Hietz and Briones 1998; Petter *et al*. 2016; Nitta *et al*. 2020).

Grouping among terrestrial species was generally less well-defined than in epiphytic species. Traits of terrestrial species were spread along a short LES-like gradient combining low Lth and high SLA and leaf N at the species and community levels. Reconstructions of the evolutionary history of functional traits across ferns lineages revealed that higher LA and mass-based leaf nitrogen are more like associated with terrestrial than epiphytes since the late Jurassic period (Jin *et al*. 2021). Although only moderate compared to the xeromorphic epiphytes, some terrestrial lineages did nonetheless express higher leaf thickness and EWT values, thus likely presenting moderate xeromorphic adaptations expressed by a subset of Athyriaceae species (*e.g., Deparia formosana*, *Diplazium donianum* var. *donianum*). In contrast, the most conservative side, including species from Plagiogyriaceae and from a subset of species from the Athyriaceae family, overlapped with the mesomorphic and drought-deciduous epiphytes in the trait space. Overall, these trait patterns support the more severe drought (only poikilohydric and pronounced xeromorphic epiphytes) and nutrient limitation (only acquisitive terrestrial species) experienced by epiphytic compared to terrestrial species at both species and community levels.

### Trait-trait relationship patterns

For terrestrial species, our analysis showed LES-related correlations at the community-level, however these relationships were weak or disappeared after correcting for phylogenetic effects at the species level. The community-level patterns agree with previous studies that observed similar LES relationships for terrestrial ferns and angiosperms (Karst and Lechowicz 2007; Campany *et al*. 2019; Lin *et al*. 2020). For epiphytic species, leaf trait patterns under the LES expectation were clearer in phylogeny corrected analysis at the species level, while at the community-level appeared to show a less expressive LES trend. The relationships between leaf N, Lth, and mass-based LA within growth habits in our study appear to be complex, nonetheless, they continue to demonstrate an overall tendency driven by general leaf economics (Onoda *et al*. 2017). Absent links between SLA × leaf N and SLA × photosynthetic rate have been observed in previous studies focusing on epiphytic ferns (Zhang *et al*. 2014; Campany *et al*. 2021). These patterns are likely due to a combination of factors. Firstly, most of epiphytic species seem to consist mostly of relatively resource-conservative species, thus lacking representatives at the acquisitive side of the spectrum, which partly reflects recent observations from trait comparisons across nearly 3000 vascular epiphytes (Hietz *et al*. 2022). Secondly, the strong divergent trait adaptations to drought across the epiphytic groups also affected traits traditionally aligned with the LES. After phylogenetic correction, LDMC was negatively related to SLA across epiphytic species at the species level; however, because of the unique trait composition of poikilohydric fern species, this trait relationship resulted in a positive correlation at the community level.

Although growth habits showed distinct trait means, there was some overlap in trait-trait correlations at both the species and community levels within species groups. This could be the result of phylogenetic constraints among in our trait – trait analysis (*e.g*., strong phylogenetic signal when traits from both growth habit were linked) or owing to coordinated responses of a few traits to environmental pressure (*e.g*., distinct traits responding similarly to the same environmental factor) (Givnish 1987). Despite some overlap in trait covariation patterns, our results show that trait patterns should not be extrapolated from terrestrial to epiphytic fern species. At the species level, each growth habit also demonstrated trait correlations that were not shared between them, which could reflect the distinct way species groups’ respond to specific environmental factors (Cardelús *et al*. 2006). This shows that trait patterns should not be extrapolated from the species to the community level, since species filtering at different sites can result in different trait-trait correlations at the community level.

### Trait-environment patterns

Despite the stark differences in mean traits between epiphytic and terrestrial species, several drought-related traits changed similarly with the climate proxies, although significant responses were observed only for terrestrial species. These responses, such as increased δ^13^C and EWT toward low elevation and fog frequency, seem to support the importance of water availability for fern community and trait composition (Kluge and Kessler 2007; Petter *et al*. 2016; Medeiros *et al*. 2019). The positive correlation of CM Lth × EWT across growth habits further illustrates their similar environmental response. The negative relationship between elevation and δ^13^C for terrestrial species suggests higher proneness to drought and associated higher water-use efficiency at low elevation (Farquhar *et al*. 1982; Maréchaux *et al*. 2020). This response was not found for epiphytic species, resulting in an unexpected negative correlation for CM δ^13^C across growth habits (Fig. 3G). Our findings might reflect the diversity lineages of epiphytic species with distinct ecologies. *Pyrrosia* species, for example, are known for their Crassulacean acid metabolism, a photosynthetic mode that conserve significant amounts of leaf water, regardless of water availability, which allow for a wider ecological niche and occurrence across different habits (Chiang *et al*. 2013).

Most of the LES traits did not respond to any of the measured climate proxies for either growth habit. Nevertheless, several studies found a significant decrease in community-level SLA with elevation for terrestrial and epiphytic ferns (Kessler *et al*. 2007; Salazar *et al*. 2012; Nitta *et al*. 2020). Furthermore, the CMs of these traits were not correlated across growth habits in our study, suggesting that different processes govern their community-level variation. Soil variation, for example, is expected to only directly impact leaf economic traits of terrestrial, but not epiphytic species (Watkins *et al*. 2007). Previous work in our study area indeed observed significant effects of soil composition on leaf N of terrestrial ferns (Helsen *et al*. 2021). LDMC showed strong relationships with elevation and fog and a moderate relationship with heat load for terrestrial species. The increased LDMC with elevation might reflect a shift to more conservative leaf traits for terrestrial species due to lower temperatures, as often observed for angiosperms (cf. Helsen *et al*. 2018), or due to decreased nutrient availability at a higher elevation. The LDMC patterns, however, more likely reflect water availability patterns. For instance, in a lower montane neotropical rainforest, Viana and Dalling (2022) found lower LDMC values of terrestrial ferns associated with aseasonal sites. The consistently negative correlation between LDMC × EWT at the species level for both growth habits further supports this. Terrestrial species showed more significant trait-environment relationships than epiphytes, potentially reflecting stronger evolutionary constraints affecting epiphytic species.

## CONCLUSIONS

Epiphytic ferns differ in mean species- and community-level trait values from terrestrial ferns along an elevation gradient in Northern Taiwan. These trait differences seem to occur because epiphytes experience higher drought and nutrient stress than terrestrial species. Trait-trait correlations traditionally associated with the LES were present at species level for terrestrial and epiphytic species. At the community level, however, these relationships were less expressive for epiphytes, likely because epiphytic trait patterns are shaped by stronger gradients in water than in nutrient availability. The LES multivariate correlations in each growth habit might represent two leaf functional traits axes of variation which in turn might be influenced by the importance of drought-stress traits. Trait– environment relationships were most prominent in terrestrial species. However, trends in relationships were similar for drought-related traits across growth habits, while LES traits were not coordinated across growth habits at the community level. Overall, these results illustrate that trait patterns are not totally equivalent for epiphytic and terrestrial species and should not be extrapolated across growth habits. Phylogenetic constraints in epiphytes should not be ruled out in explaining differences in trait responses when comparing growth habits. This study is furthermore one of the first to quantify fern functional leaf traits for the fern-rich subtropical forests of East Asia. Since no abundance data was available for the community-level trait analyses, care should be taken when comparing our patterns with studies using abundance-weighted trait means.

## SUPPLEMENTARY DATA

Supplementary data consist of the following:

Supporting_Information1: Figure S1: Map of the 59 vegetation plots in northern Taiwan; Figure S2: Climatic diagrams for precipitation and temperature from Fushan and Lalashan weather stations; Table S1: Complete species list; Appendix S1: Detailed procedures for trait measurement in ferns; Figure S3: Terminology associated with fern leaf morphology; Figure S4: Leaf locations for leaf thickness measurement; Figure S5: Leaf locations for leaf chlorophyll content measurement; Figure S6: Procedure for leaf area measurement; Appendix S2: Detailed for phylogeny reconstruction; Table S2: Results of the Mann-Whitney U tests; Table S3: PCA loadings for species level analyses; Table S4: Statistics of hypervolume analyses; Figure S7: Visualization of the posterior density of stochastic mapping for trait growth habit; Figure S8: Visualization of the trait hypervolume; Figure S9: PCA ordination for raw and independent contrasts; Figure S10: Specie-level Spearman correlations with raw trait; Figure S11: Specie-level Person correlations with independent contrasts; Table S5: Pagel’s lambda statistics; Table S6: PCA loadings for community level analyses; Figure S12: Community-level Spearman correlations; Table S7: Pairwise Spearman rank correlation of community mean trait values between growth habits.

Helsen_etal_Supporting_information_2_20230506.txt: A text file with R script for data analysis.

## ACKNOWLEDGEMENTS

We would like to thank all volunteers who contributed to fieldwork and trait measurements. TYL and DZ conceived the original idea and collected data. KH and JLV elaborated on the idea, and LYK built the phylogenetic tree. KH and JLV analysed data and wrote the manuscript with contributions from TYL and DZ. All co-authors commented on the final version of the manuscript. Data were collected as part of the Master thesis conducted by TYL at the National Taiwan University.

## DATA AVAILABILITY

Data used in this manuscript is available in https://github.com/zdealveindy/VegLab/blob/main/data/elevation-transect-ferns

