## Supporting_information_1 for "Comparing trait syndromes between Taiwanese subtropical terrestrial and epiphytic ferns at the species and community level"

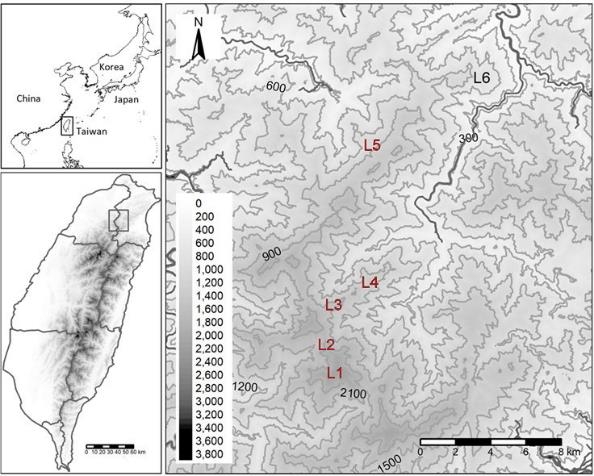

**Figure S1.** Location of the six elevation zones where 59 vegetation plots were established across northern Taiwan forests.

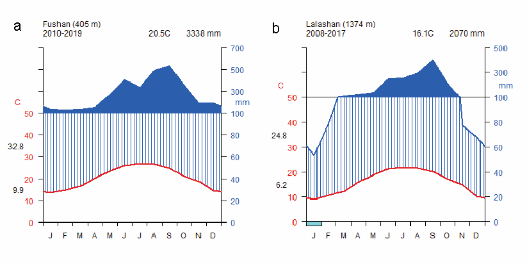

**Figure S2.** Climate diagrams for a) Fushan weather station (ID C0A560, N24.7762°, E121.5028°, 405 m a.s.l.), and b) Lalashan weather station (ID A0C54, N24.6903°, E121.4039°, 1374 m a.s.l.). Data were downloaded from the public repository of the Taiwan Central Weather Bureau (<http://e-service.cwb.gov.tw/HistoryDataQuery/>).

**Table S1.** Full fern species list from composition data. Species code provided for each terrestrial and epiphytic fern. The column ‘traits’ indicates for which species functional traits were measured. ^a^ = no leaf chlorophyll content measurement possible. Code provided for species which functional traits were collected.

| **Species** | **Traits** | **Growth habit** | **Code** |
| --- | --- | --- | --- |
| *Abacopteris gymnopteridifrons* (Hayata) Ching | × | terrestrial | Abacgymn |
| *Abrodictyum obscurum* (Blume) Ebihara & K.Iwats. |  | epiphyte |  |
| *Acystopteris taiwaniana* (Tagawa) Á.Löve & D.Löve | × | terrestrial | Acystaiw |
| *Acystopteris tenuisecta* (Blume) Tagawa |  | terrestrial |  |
| *Alsophila podophylla* Hook. | × | terrestrial | Alsopodo |
| *Alsophila spinulosa* (Wall. ex Hook.) R.M.Tryon |  | terrestrial |  |
| *Arachniodes amabilis* (Blume) Tindale var. *amabilis* | × | terrestrial | Aracamab |
| *Arachniodes festina* (Hance) Ching | × | terrestrial | Aracfest |
| *Arachniodes pseudoaristata* (Tagawa) Ohwi | × | terrestrial | Aracpseu |
| *Asplenium antiquum* Makino | × | epiphyte | Asplanti |
| *Asplenium nidus* L. | × | epiphyte | Asplnidu |
| *Asplenium normale* D.Don var. *normale* | × | terrestrial | Asplnorm |
| *Asplenium wilfordii* Mett. ex Kuhn var. *wilfordii* | × | epiphyte | Asplwilf |
| *Athyrium arisanense* (Hayata) Tagawa | × | terrestrial | Athyaris |
| *Athyrium clarkei* Baker | × | epiphyte | Athyclar |
| *Athyrium delavayi* Christ var*. delavayi* |  | terrestrial |  |
| *Athyrium iseanum* Rosenst. var*. angustisectum* Tagawa |  | terrestrial |  |
| *Athyrium nakanoi* Makino | × | terrestrial | Athynaka |
| *Athyrium opacum* (D.Don) Copel. |  | terrestrial |  |
| *Blechnopsis orientalis* (L.) C.Presl |  | terrestrial |  |
| *Cheiropleuria integrifolia* (D.C.Eaton ex Hook.) M.Kato,  Y.Yatabe, Sahashi & N.Murak. |  | terrestrial |  |
| *Coniogramme intermedia* Hieron. |  | terrestrial |  |
| *Coryphopteris angulariloba* (Ching) L.J.He & X.C.Zhang |  | terrestrial |  |
| *Coryphopteris castanea* (Tagawa) Y.H.Chang, A. Ebihara &  L.Y. Kuo | × | terrestrial | Corycast |
| *Crepidomanes minutum* (Blume) K.Iwats. subsp. *minutum* |  | epiphyte |  |
| *Davallia trichomanoides* Blume | × | epiphyte | Davatric |
| *Deparia formosana* (Rosenst.) R.Sano | × | terrestrial | Depaform |
| *Dicranopteris tetraphylla* (Rosenst.) C.M.Kuo |  | terrestrial |  |
| *Diplazium dilatatum* Blume | × | terrestrial | Dipldila |
| *Diplazium doederleinii* (Luerss.) Makino | × | terrestrial | Dipldoed |
| *Diplazium donianum* (Mett.) Tardieu var. *donianum* | × | terrestrial | Dipldoni |
| *Diplazium kawakamii* Hayata var. *kawakamii* | × | terrestrial | Diplkawa |
| *Diplazium mettenianum* (Miq.) C.Chr. | × | terrestrial | Diplmett |
| *Diplazium okinawaense* Tagawa |  | terrestrial |  |
| *Diplazium petrii* Tardieu | × | terrestrial | Diplpetr |
| *Diplazium pullingeri* (Baker) J.Sm. | × | terrestrial | Diplpull |
| *Diplazium sp.* |  | terrestrial |  |
| *Diplazium virescens* Kunze var. *virescens* |  | terrestrial |  |
| *Diplopterygium glaucum* (Thunb. ex Houtt.) Nakai | × | terrestrial | Diplglau |
| *Drynaria coronans* (Wall. ex Mett.) J.Sm. ex T.Moore |  | epiphyte |  |
| *Dryopteris formosana* (Christ) C.Chr. | × | terrestrial | Dryoform |
| *Dryopteris hasseltii* (Blume) C.Chr. | × | terrestrial | Dryohass |
| *Dryopteris hendersonii* (Bedd.) C.Chr. | × | terrestrial | Dryohend |
| *Dryopteris lepidopoda* Hayata |  | terrestrial |  |
| *Dryopteris melanocarpa* Hayata | × | terrestrial | Dryomela |
| *Dryopteris paleolata* (Pic.Serm.) Li Bing Zhang | × | terrestrial | Dryopale |

| **Species** | **Traits** | **Growth habit** | **Code** |
| --- | --- | --- | --- |
| *Dryopteris polita* Rosenst. |  | terrestrial |  |
| *Dryopteris subexaltata* (Christ) C.Chr. |  | terrestrial |  |
| *Dryopteris subtriangularis* (C.Hope) C.Chr*.* | × | terrestrial | Dryosubt |
| *Dryopteris wuzhaohongii* Li Bing Zhang | × | terrestrial | Dryowuzh |
| *Elaphoglossum yoshinagae* (Yatabe) Makino | × | epiphyte | Elapyosh |
| *Goniophlebium amoenum* (Wall. ex Mett.) Bedd. var.  *arisanense* (Hayata) Rödl-Linder | × | epiphyte | Goniaris |
| *Goniophlebium mengtzeense* (Christ) Rödl-Linder | × | epiphyte | Gonimeng |
| *Goniophlebium raishaense* (Rosenst.) L.Y.Kuo |  | epiphyte |  |
| *Haplopteris flexuosa* (Fée) E.H.Crane | × | epiphyte | Haplflex |
| *Histiopteris incisa* (Thunb.) J.Sm. | × | terrestrial | Histinci |
| *Hymenasplenium adiantifrons* (Hayata) Viane & S.Y.Dong | × | terrestrial | Hymeadia |
| *Hymenophyllum badium* Hook. & Grev. | × | epiphyte | Hymebadi |
| *Hymenophyllum nitidulum* (Bosch) Ebihara & K.Iwats. | ×a | epiphyte | Hymeniti |
| *Hymenophyllum okadae* Masam. | × | epiphyte | Hymeokad |
| *Hymenophyllum oligosorum* Makino | × | epiphyte | Hymeolig |
| *Hymenophyllum paniculiflorum* C.Presl | ×a | epiphyte | Hymepani |
| *Hymenophyllum polyanthos* (Sw.) Sw. | × | epiphyte | Hymepoly |
| *Lemmaphyllum microphyllum* C.Presl | × | epiphyte | Lemmmicr |
| *Lepidomicrosorium ningpoense* (Baker) L.Y.Kuo | × | epiphyte | Lepining |
| *Lepisorus kawakamii* (Hayata) Tagawa | × | epiphyte | Lepikawa |
| *Lepisorus monilisorus* (Hayata) Tagawa | × | epiphyte | Lepimoni |
| *Lepisorus obscurevenulosus* (Hayata) Ching | × | epiphyte | Lepiobsc |
| *Lepisorus rostratus* (Bedd.) Tagawa | × | epiphyte | Lepirost |
| *Lepisorus suboligolepidus* Ching | × | epiphyte | Lepisubo |
| *Leptogramma tottoides* Hayata ex H.Ito |  | terrestrial |  |
| *Lindsaea bonii* Christ | × | terrestrial | Lindboni |
| *Lindsaea chienii* Ching | × | terrestrial | Lindchie |
| *Loxogramme remotefrondigera* Hayata | × | epiphyte | Loxoremo |
| *Loxogramme salicifolia* (Makino) Makino | × | epiphyte | Loxosali |
| *Metathelypteris gracilescens* (Blume) Ching | × | terrestrial | Metagrac |
| *Metathelypteris laxa* (Franch. & Sav.) Ching |  | terrestrial |  |
| *Metathelypteris uraiensis* (Rosenst.) Ching | × | terrestrial | Metaurai |
| *Microlepia hookeriana* (Wall. ex Hook.) C.Presl | × | terrestrial | Micrhook |
| *Microlepia obtusiloba* Hayata |  | terrestrial |  |
| *Micropolypodium okuboi* (Yatabe) Hayata | ×a | epiphyte | Microkub |
| *Monachosorum henryi* Christ | × | terrestrial | Monahenr |
| *Monachosorum maximowiczii* (Baker) Hayata |  | epiphyte |  |
| *Nephrolepis cordifolia* (L.) C.Presl |  | terrestrial |  |
| *Odontosoria chinensis* (L.) J.Sm. |  | terrestrial |  |
| *Plagiogyria adnata* (Blume) Bedd. | × | terrestrial | Plagadna |
| *Plagiogyria euphlebia* (Kunze) Mett. | × | terrestrial | Plageuph |
| *Plagiogyria falcata* Copel. | × | terrestrial | Plagfalc |
| *Plagiogyria glauca* (Blume) Mett. | × | terrestrial | Plagglau |
| *Plagiogyria stenoptera* (Hance) Diels | × | terrestrial | Plagsten |
| *Polystichum hancockii* (Hance) Diels | × | terrestrial | Polyhanc |
| *Polystichum integripinnum* Hayata | × | terrestrial | Polyinte |
| *Polystichum parvipinnulum* Tagawa | × | terrestrial | Polyparv |
| *Prosaptia formosana* (Hayata) T.C.Hsu | × | epiphyte | Prosform |
| *Pteris bella* Tagawa | × | terrestrial | Pterbell |
| *Pteris tokioi* Masam. | × | terrestrial | Ptertoki |
| *Pteris wallichiana* J.Agardh |  | terrestrial |  |
| *Pyrrosia lingua* (Thunb.) Farw. | × | epiphyte | Pyrrling |

| **Species** | **Traits** | **Growth habit** | **Code** |
| --- | --- | --- | --- |
| *Pyrrosia polydactyla* (Hance) Ching | × | epiphyte | Pyrrpoly |
| *Pyrrosia sheareri* (Baker) Ching | × | epiphyte | Pyrrshea |
| *Selliguea engleri* (Luerss.) Fraser-Jenk. | × | epiphyte | Sellechi |
| *Selliguea lehmannii* (Mett.) Ching | × | epiphyte | Sellengl |
| *Sitobolium zeylanicum* (Sw.) L.A. Triana & Sundue | × | terrestrial | Sitozeyl |
| *Stegnogramma griffithii* (T.Moore) K.Iwats. | × | terrestrial | Steggrif |
| *Stegnogramma wilfordii* (Hook.) Seriz*.* | × | terrestrial | Stegwilf |
| *Vandenboschia auriculata* (Blume) Copel. | × | epiphyte | Vandauri |
| *Vandenboschia kalamocarpa* (Hayata) Ebihara | ×a | epiphyte | Vandkala |
| *Woodwardia unigemmata* (Makino) Nakai |  | terrestrial |  |

**Appendix S1.** Extended trait measurement protocol for fern leaves.

### Leaf thickness

Before trait measurements, stipes were removed for all the selected leaf samples (for the terminology of fern leaf morphology, see Fig. S3). Leaf thickness (Lth, mm) was measured with a 1 µm resolution digital thickness gauge (DML3034, Digital Micrometers Ltd., UK). Leaf thickness was measured four times, at the upper left, upper right, lower left and lower right of the leaf (i.e. lamina), and averaged for each leaf. For simple laminas, the four measurement points were positioned at an equal distance from the midrib and the leaf margin (Fig. S4). For ferns with compound laminas, the four measurement points were positioned in the middle of the pinnae, pinnules and pinnulets for single, double and triple-compound leaves, respectively, at an equal distance from the costa and the margin. Care was taken to avoid obvious veins and sori. When this proved difficult, part of the lamina was cut to ease leaf thickness measurement. Note that any cut-off segments was carefully preserved for leaf area measurement (see further).

### Area-based leaf chlorophyll content

Area-based chlorophyll content (Chl) was measured using a Soil Plant Analysis Development (SPAD-502) chlorophyll meter (Konica Minolta, Japan). For each leaf, we performed one to twenty measurements on each side (left and right) of the lamina. For compound lamina with many pinnae, chlorophyll was measured every two or three pinnae. The number of measurements depended on the size of the lamina and the number of the pinnae (Fig. S5).

### Leaf area

Leaf area (LA, cm^2^) was measured on each rehydrated leaf using a scanner (Perfection V370 PHOTO & Perfection V600 PHOTO, EPSON, Japan) and the Image J software (Rueden et al. 2017). Before scanning, laminas were dissected to remove the free axes and prevent overlapping segments. Simple, palmatifid, pinnatifid, pinnatisect and bipinnatifid laminas were scanned directly. When lobes or pinnae overlapped, we cut along the rachis to separate them. For pinnate and pinnate-pinnatifid laminas, we first removed the full rachis and scanned the pinnae separately (Fig. S6). For bipinnate and bipinnate-pinnatifid, tripinnate and higher-order compound laminas, the rachis and costas were first removed before scanning the pinnules separately.

For bipinnate, bipinnatifid, tripinnate or more divided ferns with very fine segments, only a subset of all pinnae of the lamina were dissected (one every two, three or four pinnae pairs) as described earlier for measuring SLA, LDMC, and EWT (see further) to save time. All remaining pinnae of the lamina were only dissected by cutting the costa to prevent overlapping of the pinnules and were then scanned for calculating leaf area.

### Specific leaf area, leaf dry matter content and equivalent water thickness

Right after scanning, laminas (or the dissected pinnae or pinnules) were weighed using a 0.1 mg precision balance (Adventurer AR2140, OHAUS Corp. USA or XS 225A-SCS, Precisa, Switzerland) to obtain leaf fresh weight (g). All lamina, pinnae and pinnule parts were then transferred to one labelled paper bag per original leaf and oven dried at 70°C for at least 72 hours. After drying, dry weight (mg) was measured with the precision balance. Specific leaf area (SLA, mm^2^/mg) was then calculated by dividing the one-sided leaf area by the leaf dry weight. Leaf dry matter content (LDMC, mg/g) was calculated as the leaf dry weight divided by the leaf fresh weight. Equivalent water thickness (EWT, mg/mm^2^), which quantifies area- based water content was calculated as (leaf fresh weight – leaf dry weight)/ leaf area.

### Leaf nitrogen content and leaf ^13^C/^12^C and ^15^N/^14^N stable isotope ratios.

After measuring leaf dry weights, one to four dried leaf samples of each species were selected to quantify leaf nitrogen content (leaf N, mg/g), leaf ^13^C/^12^C stable isotope ratio (δ^13^C,

‰) and leaf ^15^N/^14^N stable isotope ratio (δ^15^N, ‰). These leaves were selected based on the following criteria: 1) Preferably select a leaf from an individual with sporophylls; 2) select the leaves with the highest dry weight; 3) select leaves from plots closest to the mountain ridge (to minimize potential effects of plot aspect); 4) if for a given individual, both fertile and sterile leaves are available, both are selected; 5) if the species occurred across several elevation zones, leaves were chosen to reflect its full elevation range. The selected samples were then ground into a fine powder with the help of scissors, a mortar and a pestle. 2.000 ± 0.100 mg of the leaf powder was then weighed and placed into a tin capsule. Leaf N was measured on the tin capsules with a FlashEA 1112 series elemental analyzer (Thermo Fisher Scientific, Italy), while δ^13^C and δ^15^N were subsequently measured on the same sample with a Delta V Advantage isotope ratio mass spectrometer (Finnigan Mat, Germany), using Peedee belemnite (PDB) and atmospheric nitrogen as global standards for δ^13^C and δ^15^N, respectively.

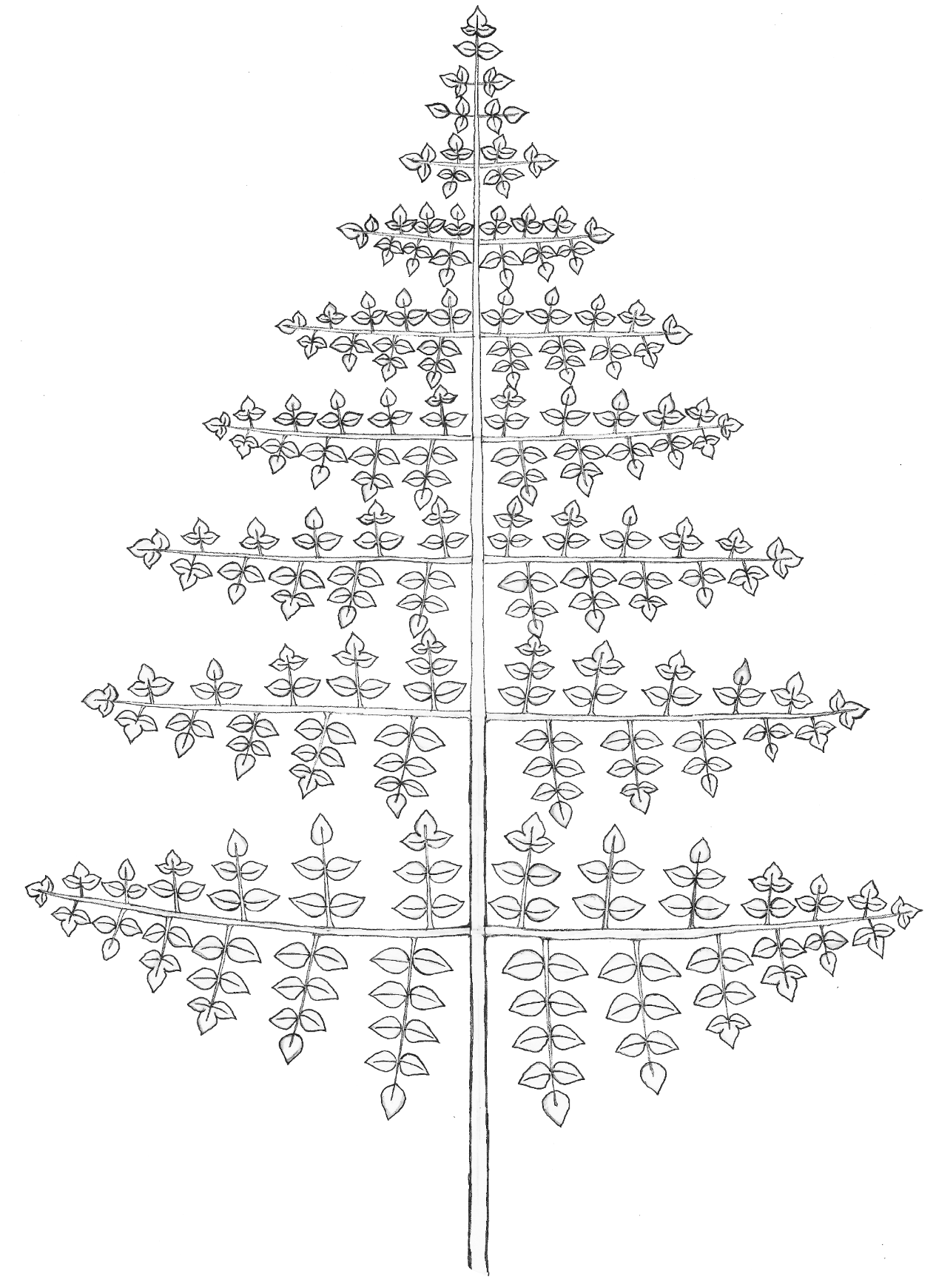

**2**

**6**

**7**

**5**

**4**

**3**

**1**

**Figure S3.** Terminology associated with fern leaf morphology. Axes terminology: 1. stipe, 2. rachis, 3. costa, 4. costule. Lamina structure terminology: 5. pinna, 6. pinnule, 7. pinnulet.

a	b	c

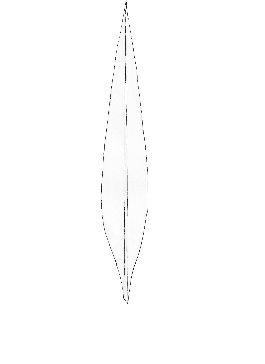

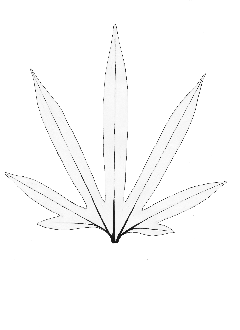

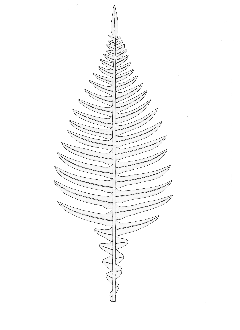

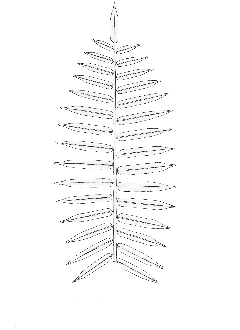

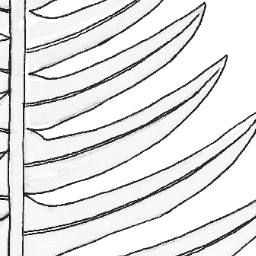

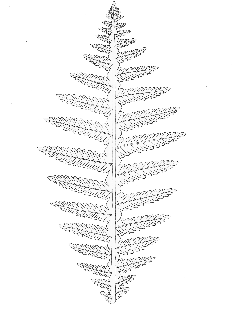

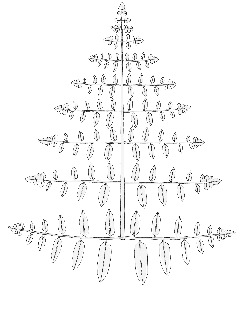

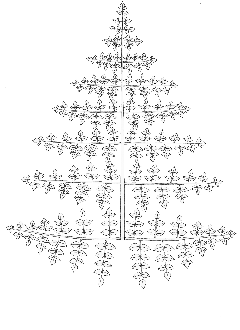

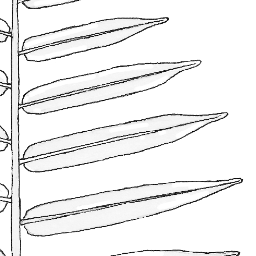

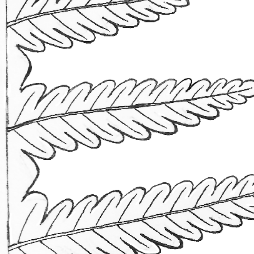

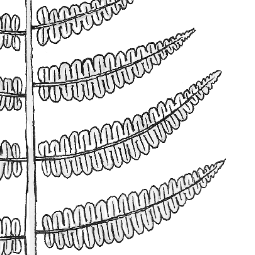

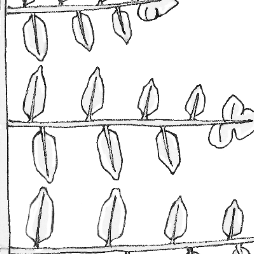

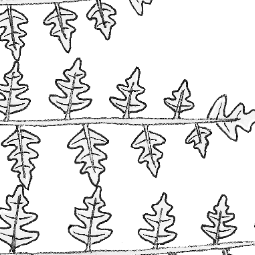

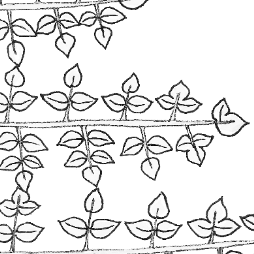

d

e

f

g

h

i

j

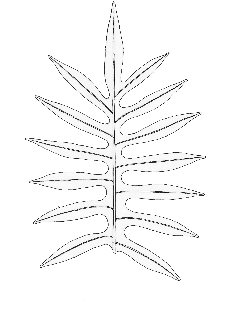

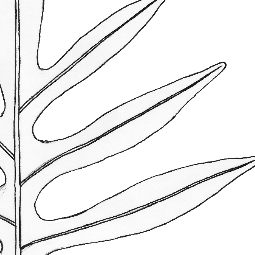

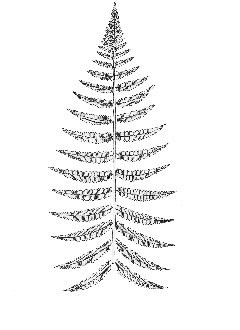

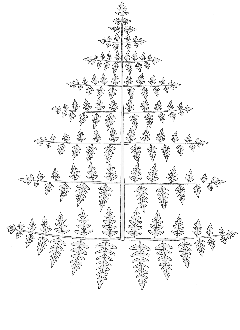

**Figure S4.** The positions for measuring leaf thickness for a) simple, b) palmatifid, c) pinnatisect, d) pinnatifid, e) pinnate, f) bipinnatifid, g) pinnate-pinnatifid, h) bipinnate, i) bipinnate-pinnatifid, and j) tripinnate laminae.

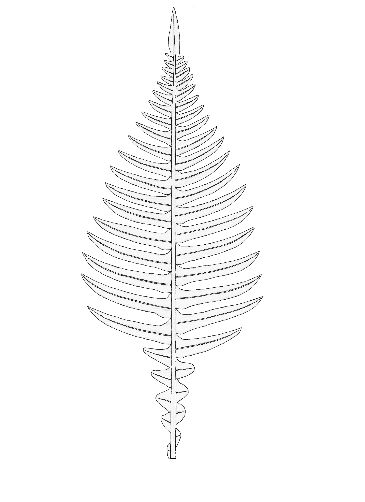

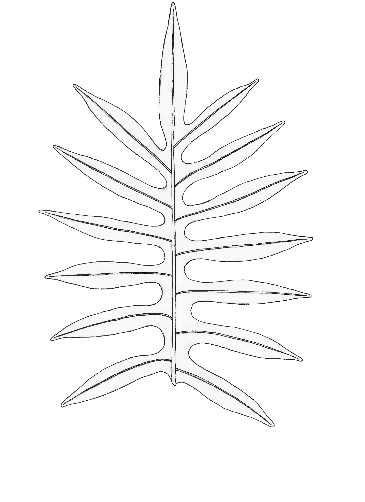

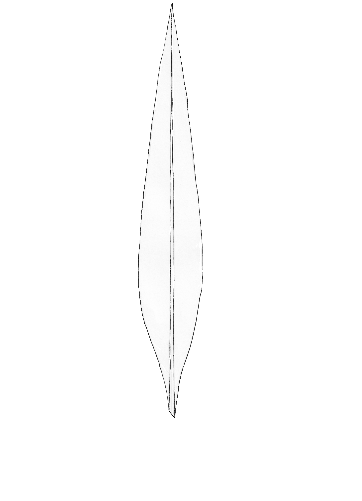

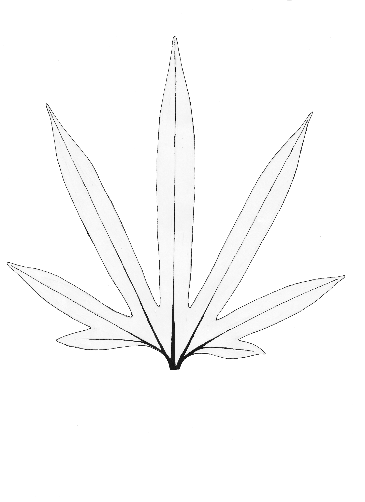

a

b

c

d

e	f	g

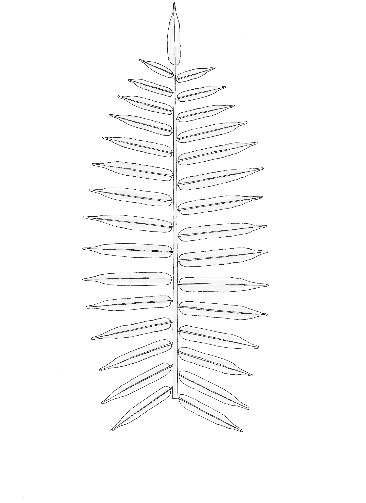

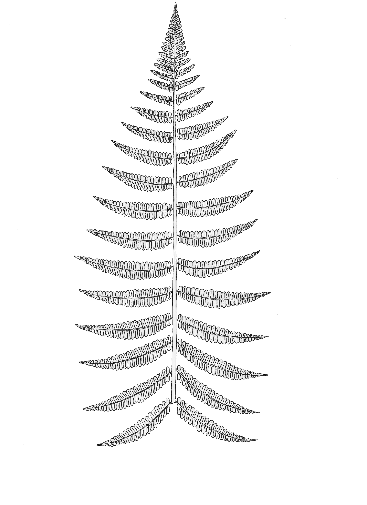

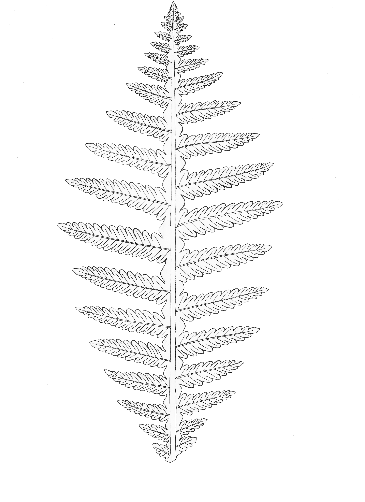

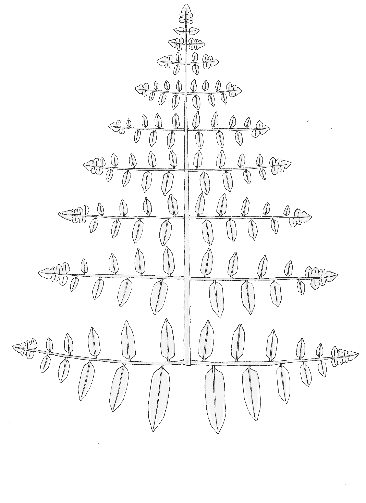

h

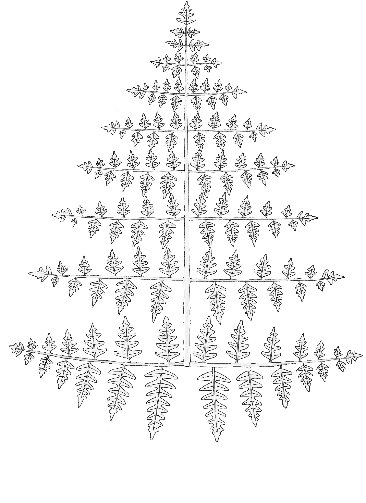

i

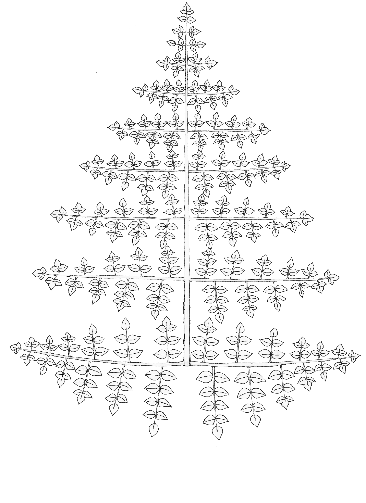

j

**Figure S5.** The positions for measuring leaf chlorophyll content for a) simple, b) palmatifid, c) pinnatisect, d) pinnatifid, e) pinnate, f) bipinnatifid, g) pinnate-pinnatifid, h) bipinnate, i) bipinnate-pinnatifid and j) tripinnate laminae.

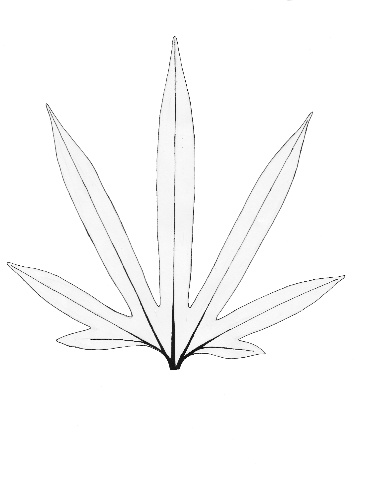

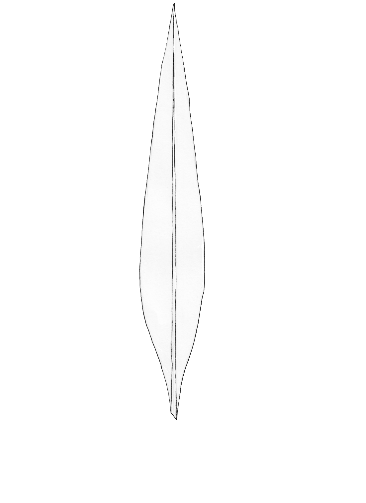

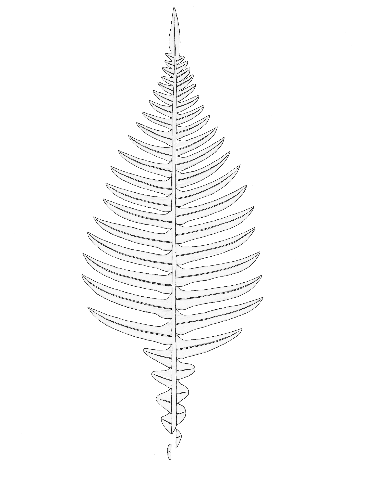

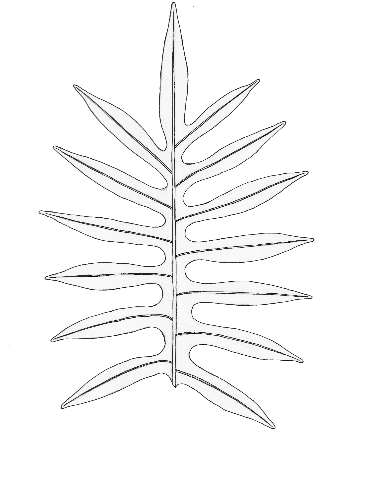

a

b

c

d

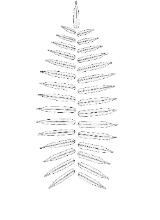

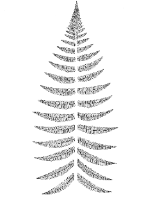

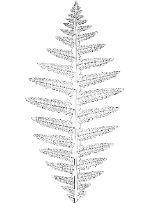
e	f	g

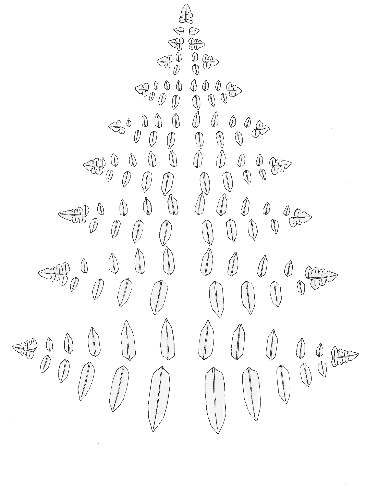

h

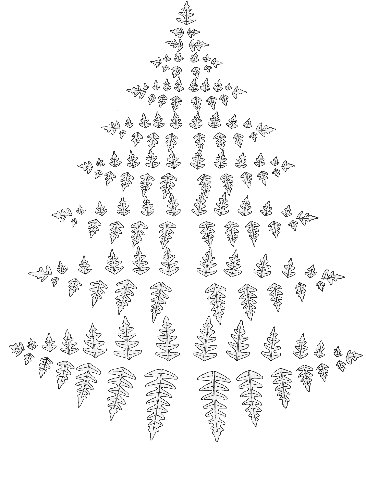

i

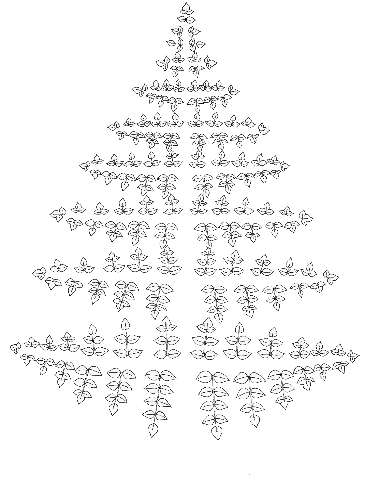

j

**Figure S6.** Procedure of leaf area measurement of a) simple, b) palmatifid, c) pinnatisect, d) pinnatifid, e) pinnate, f) bipinnatifid, g) pinnate-pinnatifid, h) bipinnate, i) bipinnate- pinnatifid and j) tripinnate laminae.

**Appendix S2.** Method for phylogenetic tree estimation using rbcL nucleotide sequences from fern species.

To infer a phylogenetic tree for our taxa, we first determined which species from our study were not represented in the dated fern phylogeny generated by Nitta *et al.* (2022). For those species (ten species: *Hymenasplenium adiantifrons, Hymenophyllum okadae, Lepidomicrosorium ningpoense, Loxogramme remotefrondigera, Plagiogyria falcata, Polystichum integripinnum, Prosaptia formosana, Pteris tokioi, Selliguea echinospora, Vandenboschia kalamocarpa*), *rbcL* were obtained fom the GBIF dataset of Asian fern DNA barcodes (Chen and Kuo 2023; https://doi.org/10.15468/nhsnn2) or NCBI GenBank (accession numbers: AB574845 and AB196364) and combined with those from FTOL v1.1.0 (Fern Tree of Life, Nitta *et al.* 2022). Sequence alignment was done using the MUSCLE algorithm (Edgar 2004) in AliView (Larsson 2014). Tree estimation was performed using maximum likelihood methods (ML) in IQ-TREE (Minh *et al.* 2020), with GTR + F + I + G4 as substitution models. Then, the final ML tree was dated using the same fossil setting as FTOL v1.1.0 (Nitta *et al.* 2022) with treePL (Smith and O’Meara 2012). The main big tree was pruned to keep only the species for downstream analysis. We calculated independent contrast for each of the nine continuous traits for all species combined using the ‘pic’ function from the *ape* R package (Paradis and Schliep 2019), under the assumption of the Brownian motion model of evolution (Felsenstein 1985). Independent contrasts were used to perform Pearson correlations and principal components analyses.

**Table S2.** Results of tests for differences in trait values between epiphytic and terrestrial species (Mann-Whitney U test). At the species level, differences between epiphytic and terrestrial species were calculated using raw species traits. At the community level, differences were tested using trait means. Test statistics (*U*) and p-value (after FDR correction, *P*-adj.) were provided. Significant *U* statistics are in bold. ^L^ = Logarithmically transformed variables for both epiphytic and terrestrial species.

| **Traits** | **Species level** | | **Community level** | |
| --- | --- | --- | --- | --- |
|  | *U* | *P*-adj. | *U* | *P*-adj. |
| SLA | **428** | < 0.001 | **490** | < 0.001 |
| LDMC | 927 | 0. 2517 | **2390** | < 0.001 |
| Leaf area^L^ | **301** | < 0.001 | **0** | < 0.001 |
| Leaf thickness^L^ | 1014 | 0.0509 | 2057 | 0.088 |
| Chlorophyll content | 721 | 0.4582 | **400** | < 0.001 |
| EWT^L^ | **1047** | 0.0259 | **2811** | < 0.001 |
| δ13C | **1224** | < 0.001 | **3365** | < 0.001 |
| δ15N | **316** | < 0.001 | **0** | < 0.001 |
| Leaf N | **133** | < 0.001 | **0** | < 0.001 |

**Table S3.** Loadings and cumulative variation for the first three principal component axes (PC) for the three ordinations on all species combined, epiphytic species and terrestrial species. Main contributors to each ordination axis (with an absolute value of loading higher than 0.30) are in bold. δ^13^C = the leaf ^13^C/^12^C stable isotope ratio, δ^15^N = the leaf ^15^N/^14^N stable isotope ratio*,* EWT = equivalent water thickness*.* ^L^ = Logarithmically transformed variables for both epiphytic and terrestrial species. A) raw trait data; B) independent contrasts.

| A | | | | | | | | | |
| --- | --- | --- | --- | --- | --- | --- | --- | --- | --- |
|  | All species raw trait data | | | Epiphytic species raw trait data | | | Terrestrial species raw trait data | | |
| traits | PC1 | PC2 | PC3 | PC1 | PC2 | PC3 | PC1 | PC2 | PC3 |
| SLA | **0.46** | -0.18 | -0.24 | **0.45** | -0.09 | -0.01 | **0.50** | -0.05 | -0.19 |
| LDMC | 0.29 | **0.38** | **0.45** | **0.39** | 0.25 | 0.12 | -0.16 | **0.63** | 0.11 |
| Leaf area^L^ | 0.02 | **-0.51** | **0.41** | -0.06 | **-0.58** | **0.32** | -0.21 | 0.17 | **-0.52** |
| Leaf thickness^L^ | **-0.49** | -0.12 | -0.10 | **-0.46** | -0.06 | -0.06 | **-0.38** | **-0.38** | 0.17 |
| Chlorophyll content | **-0.40** | -0.19 | 0.20 | **-0.40** | -0.03 | -0.12 | **-0.46** | 0.01 | 0.03 |
| EWT^L^ | **-0.50** | -0.03 | -0.18 | **-0.47** | -0.02 | -0.03 | **-0.30** | **-0.55** | 0.01 |
| δ13C | -0.13 | 0.27 | **0.48** | -0.05 | -0.17 | **0.84** | -0.21 | 0.07 | **-0.57** |
| δ15N | 0.05 | **-0.45** | **0.47** | 0.07 | **-0.56** | **-0.31** | **-0.34** | 0.13 | -0.28 |
| Leaf nitrogen | 0.22 | **-0.49** | -0.19 | 0.22 | **-0.50** | -0.23 | 0.27 | **-0.33** | **-0.50** |
| Cumul. variation (%) | 41.49 | 65.72 | 78.65 | 49.25 | 68.45 | 81.15 | 40.23 | 62.70 | 85.21 |
| B | | | | | | | | | |
|  | All independent contrasts | | | Epiphytic independent contrasts | | | Terrestrial independent contrasts | | |
| traits | PC1 | PC2 | PC3 | PC1 | PC2 | PC3 | PC1 | PC2 | PC3 |
| SLA | **0.48** | -0.28 | 0.16 | **0.53** | -0.07 | 0.28 | **0.43** | **-0.34** | 0.06 |
| LDMC | 0.17 | **0.65** | -0.15 | -0.16 | **-0.45** | **-0.44** | 0.27 | **0.52** | -0.08 |
| Leaf area^L^ | 0.21 | 0.14 | 0.20 | 0.20 | -0.09 | -0.28 | 0.20 | -0.02 | **-0.70** |
| Leaf thickness^L^ | **-0.49** | -0.24 | 0.03 | **-0.35** | **0.39** | 0.21 | **-0.49** | -0.13 | 0.08 |
| Chlorophyll content | **-0.38** | **0.35** | 0.16 | **-0.40** | -0.26 | **-0.33** | **-0.42** | 0.16 | -0.21 |
| EWT^L^ | **-0.51** | -0.22 | 0.01 | **-0.40** | **0.39** | 0.14 | **-0.50** | -0.11 | 0.02 |
| δ13C | -0.12 | **0.45** | 0.17 | **-0.33** | **-0.31** | 0.18 | -0.05 | **0.40** | -0.22 |
| δ15N | -0.16 | 0.16 | **0.60** | **-0.31** | **-0.30** | **0.43** | -0.14 | -0.21 | **-0.62** |
| Leaf nitrogen | 0.14 | -0.16 | **0.71** | 0.05 | **-0.47** | **0.51** | 0.09 | **-0.60** | -0.10 |
| Cumul. variation (%) | 35.13 | 54.30 | 69.43 | 32.03 | 58.65 | 71.79 | 40.05 | 60.84 | 74.32 |

**Table S4.** Hypervolume size and overlap statistics. Average (µ) given for the distance between hypervolume centroid, minimum, Jaccard similarity between hypervolumes, and the unique volume fraction of the hypervolumes. The unit of hypervolume sizes is expressed in standard deviation to the power of the number of trait dimensions used (SD^3^ ^–^ the first three PCA axes generated with standardized values of nine measured traits). Species-level hypervolume statistics were calculated with raw trait data and community-level hypervolume statistics with community mean trait values. Vol. = volume.

|  | ***Hypervolume size*** |  |
| --- | --- | --- |
|  | Vol. of terrestrials | 0.055 |
|  | Vol. of epiphytes | 0.081 |
|  | Vol. of the intersection | 0.021 |
|  | Vol. of the union | 0.115 |
| **Species level** | ***Distance between volumes*** |  |
|  | Centroid (µ) | 0.186 |
|  | Minimum (µ) | 0.004 |
|  | ***Overlap statistics*** |  |
|  | Jaccard similarity (µ) | 0.191 |
|  | Unique vol. fraction of terrestrials | 0.034 |
|  | Unique vol. fraction of epiphytes | 0.058 |
|  | Unique fraction terrestrials (%) | 61.62 |
|  | Unique fraction epiphytes (%) | 72.94 |
|  | ***Hypervolume size*** |  |
|  | Vol. of terrestrials | 0.018 |
|  | Vol. of epiphytes | 0.031 |
|  | Vol. of the intersection | 0.003 |
| **Community level** | Vol. of the union | 0.046 |
|  | ***Distance between volumes*** |  |
|  | Centroid (µ) | 0.197 |
|  | Minimum (µ) | 0.003 |
|  | ***Overlap statistics*** |  |
|  | Jaccard similarity (µ) | 0.044 |
|  | Unique vol. fraction of terrestrials | 0.015 |
|  | Unique vol. fraction of epiphytes | 0.028 |
|  | Unique fraction terrestrials (%) | 88.48 |
|  | Unique fraction epiphytes (%) | 93.15 |

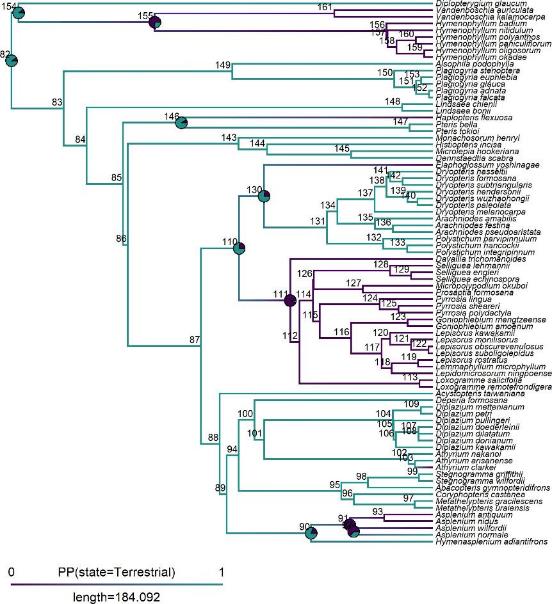

**Figure S7.** Maximum likelihood phylogenetic tree for all species combined showing the posterior density of stochastic mapping for trait growth habit. Selected nodes (10 nodes) displaying a pie chart showed the probability of epiphytic state (0) or terrestrial state (1). Numbers are node labels, green branches and nodes are terrestrial state, purple are epiphytic state.

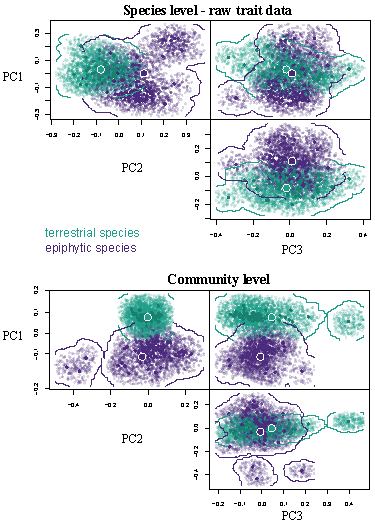

**Figure S8.** Visualization of the trait hypervolume in the first three PCA dimensions calculated on all species combined using raw trait data for species-level and trait means for community- level analyses. Large points depict hypervolume centroids, medium points depict data points (species × traits combinations in case of the species-level analysis), and small points depict randomized points sampled from the inferred hypervolume to visualize the stochastic description of each hypervolume for species and community levels respectively.

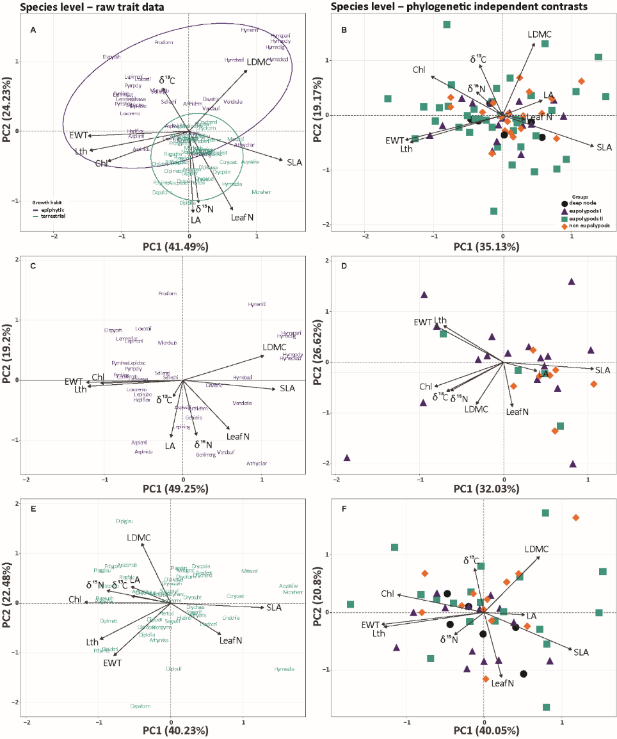

**Figure S9.** Biplots for the principal component analyses, to compare analysis done on raw trait data (left column) with the analysis done on phylogenetic independent contrasts (right column).

A. the full species × trait matrix, B. the phylogenetic independent contrasts × trait matrix, C. epiphytic species × trait matrix, D. phylogenetic independent contrasts associated with epiphytic growth habit × epiphytic trait matrix, E. terrestrial species × trait matrix, and F. phylogenetic independent contrasts associated with terrestrial growth habit × terrestrial trait matrix. Chl = leaf chlorophyll content, δ^13^C *=* the leaf ^13^C/^12^C stable isotope ratio, δ^15^N *=* the leaf ^15^N/^14^N stable isotope ratio*,* EWT = equivalent water thickness, Lth = leaf thickness, LA = leaf area, Leaf N = leaf nitrogen content, and LDMC = leaf dry matter content. Deep nodes = nodes on the dated tree not associated with groups defined in this study (eupolypods I, eupolypods II, non eupolypods).

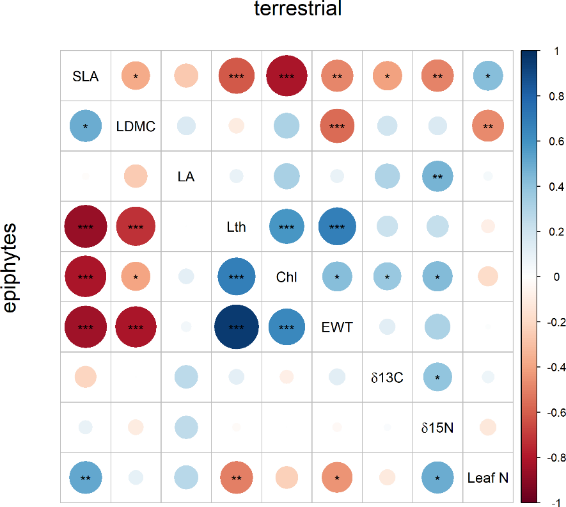

**Figure S10.** Species-level trait-trait correlations within species groups. Pairwise Spearman rank correlations were calculated for each trait using **raw traits**. Top triangle is for terrestrial species. The bottom triangle is for epiphytic species. Colours and size relate to the strength of correlation (ρ). Significant correlations: * = p < 0.05, ** = p < 0.01, and *** p < 0.001. SLA = specific leaf area, LDMC = leaf dry matter content, Lth = leaf thickness, Chl = leaf chlorophyll content, EWT = equivalent water thickness, δ^13^C = the leaf ^13^C/^12^C stable isotope ratio, δ^15^N = the leaf ^15^N/^14^N stable isotope ratio., leaf N = leaf nitrogen.

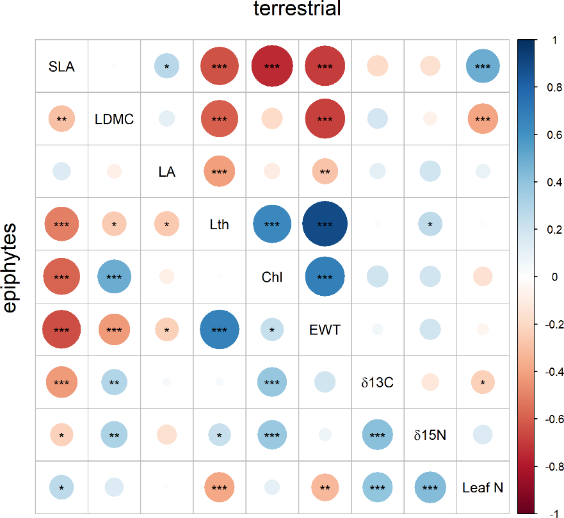

**Figure S11.** Species-level trait-trait correlations within species groups. Pairwise Pearson correlations were calculated for each trait using **independent contrasts**. Top triangle is for terrestrial species. The bottom triangle is for epiphytic species. Colours and size relate to the strength of correlation (ρ). Significant correlations: * = p < 0.05, ** = p < 0.01, and *** p <

0.001. SLA = specific leaf area, LDMC = leaf dry matter content, Lth = leaf thickness, Chl = leaf chlorophyll content, EWT = equivalent water thickness, δ^13^C = the leaf ^13^C/^12^C stable isotope ratio, δ^15^N = the leaf ^15^N/^14^N stable isotope ratio., leaf N = leaf nitrogen.

**Table S5.** Pagel’s λ statistic and probabilities for each continuous trait for all species combined. Lambda (λ) = fitted value of *λ*; logL = log likelihood for fitted Lamba; logL0 = log likelihood for Lambda equals 0; *P =* p-values; *P*-adj. = p-values adjusted with the false discovery rate method; ^L^ = log transformed.

| **trait** | **Lambda (λ)** | ***logL*** | ***logL0*** | ***P*** | ***P*-adj.** |
| --- | --- | --- | --- | --- | --- |
| SLA | 0.97 | -298.00 | -322.00 | < 0.001 | < 0.001 |
| LDMC | 0.78 | -447.00 | -470.00 | < 0.001 | < 0.001 |
| Leaf area ^L^ | 0.91 | -69.30 | -92.30 | < 0.001 | < 0.001 |
| Leaf thickness ^L^ | 0.92 | -5.37 | -39.40 | < 0.001 | < 0.001 |
| Chlorophyll content | 0.87 | -278.00 | -300.00 | < 0.001 | < 0.001 |
| EWT ^L^ | 0.98 | 9.12 | -24.00 | < 0.001 | < 0.001 |
| δ13C | 0.60 | -156.00 | -161.00 | 0.0027 | 0.024 |
| δ15N | 0.55 | -120.00 | -128.00 | < 0.001 | < 0.001 |
| Leaf N | 0.89 | -256.00 | -276.00 | < 0.001 | < 0.001 |

**Table S6.** Trait loadings and cumulative variation for the first three principal component axes (PC) for the three plot × community mean (CM) trait ordinations, for all species combined, only for epiphytic species and for terrestrial species. Main contributors to each ordination axis (with an absolute value of loading higher than 0.30) are in bold. δ^13^C = the leaf ^13^C/^12^C stable isotope ratio, δ^15^N = the leaf ^15^N/^14^N stable isotope ratio*,* EWT = equivalent water thickness*.* ^L^

= log transformed.

| for all species | | | | epiphytic species | | | terrestrial species | | |
| --- | --- | --- | --- | --- | --- | --- | --- | --- | --- |
|  | PC1 | PC2 | PC3 | PC1 | PC2 | PC3 | PC1 | PC2 | PC3 |
| CM SLA | 0.26 | **-0.39** | **-0.49** | **0.48** | -0.03 | -0.10 | **0.39** | -0.26 | 0.04 |
| CM LDMC | -0.21 | **-0.42** | **0.53** | -0.27 | -**0.52** | 0.06 | **0.41** | -0.11 | 0.11 |
| CM Leaf area ^L^ | **0.46** | 0.05 | 0.14 | -0.18 | 0.13 | -**0.87** | **-0.31** | -0.28 | 0.29 |
| CM Leaf thickness ^L^ | 0.02 | **0.54** | -0.14 | -0.17 | **0.57** | 0.25 | **-0.42** | 0.21 | -0.10 |
| CM Chlorophyll content | **0.33** | **0.30** | **0.36** | **-0.44** | 0.01 | -0.07 | **-0.36** | -0.23 | **-0.35** |
| CM EWT ^L^ | -0.14 | **0.52** | -0.14 | -0.21 | **0.56** | -0.01 | **-0.41** | 0.23 | -0.13 |
| CM δ^13^C | **-0.42** | 0.10 | 0.21 | -0.28 | -0.22 | 0.09 | -0.03 | **0.52** | **0.68** |
| CM δ^15^N | **0.41** | 0.07 | **0.47** | **-0.44** | -0.14 | -0.15 | -0.29 | -0.29 | **0.48** |
| CM Leaf nitrogen | **0.46** | -0.07 | -0.14 | **0.34** | -0.02 | -**0.36** | -0.15 | **-0.58** | 0.24 |
| Cumul. variation (%) | 48.01 | 84.5 | 93.31 | 45.98 | 72.33 | 83.3 | 56.39 | 78.65 | 88.36 |

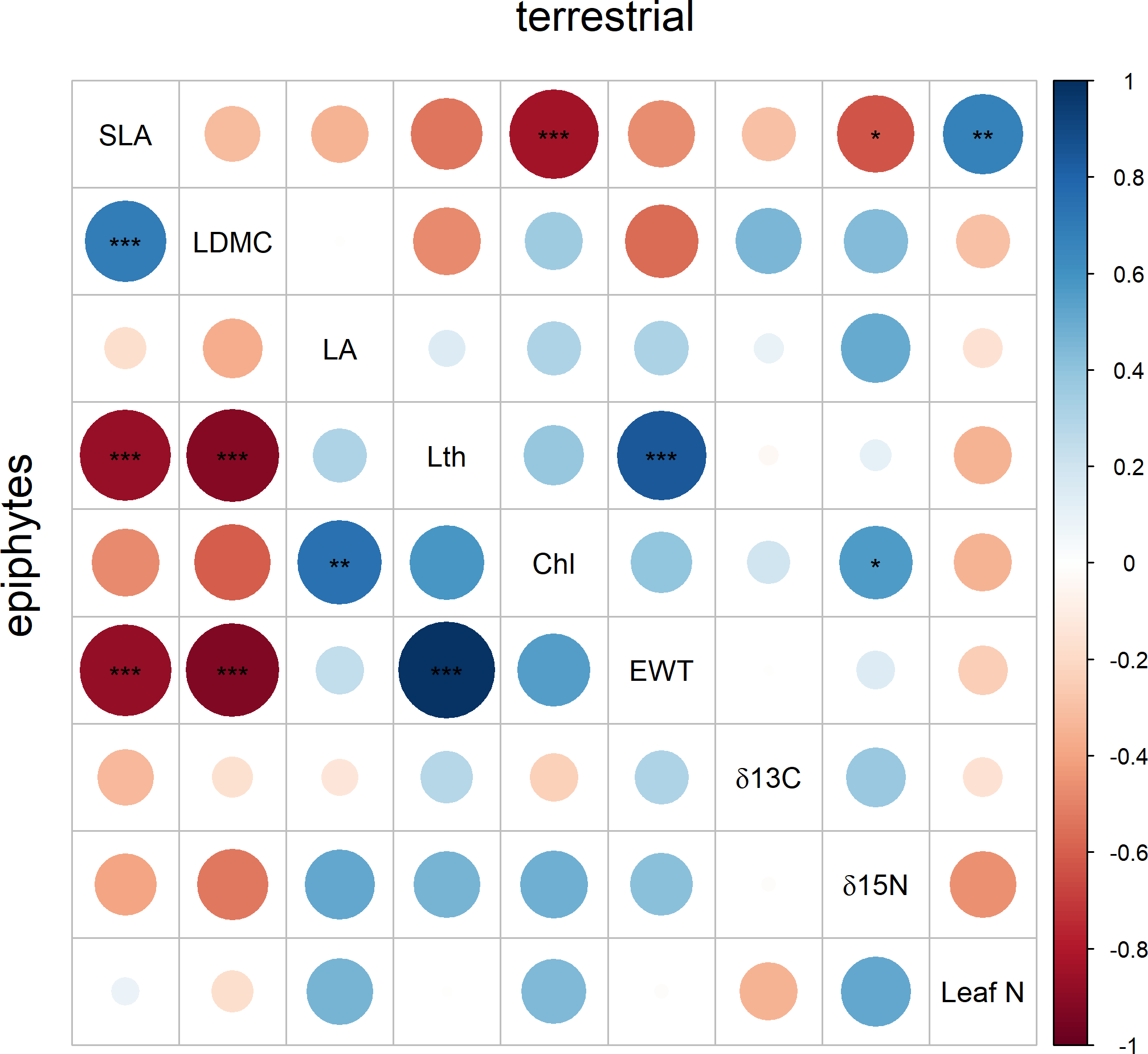

**Figure S12.** Community-level trait-trait correlations within species groups. Pairwise Spearman rank correlations between all measured **CM traits at the community level**. The top triangle is for terrestrial species. The bottom triangle is for epiphytic species. Colours relate to the strength of correlation (ρ). Significant correlations: * = p < 0.05, ** = p < 0.01, and *** = p < 0.001. SLA = specific leaf area, LDMC = leaf dry matter content, Lth = leaf thickness, Chl = leaf chlorophyll content, EWT = equivalent water thickness, δ^13^C = the leaf ^13^C/^12^C stable isotope ratio, δ^15^N = the leaf ^15^N/^14^N stable isotope ratio.

**Table S7**. Pairwise Spearman rank correlation of community mean trait values between the epiphytic and terrestrial species. ρs statistic and p-value (after FDR correction, *P*-adj.) given. Significant results are in bold. δ^13^C = the leaf 13C/12C stable isotope ratio, δ^15^N = the leaf 15N/14N stable isotope ratio, EWT = equivalent water thickness, leaf N = leaf nitrogen. ^L^ Logarithmic transformation for both epiphytic and terrestrial species.

| **Traits** | **ρs** | ***P*-adj.** |
| --- | --- | --- |
| SLA | 0.13 | 0.372 |
| LDMC | 0.27 | 0.091 |
| Leaf area ^L^ | -0.15 | 0.344 |
| **Leaf thickness** ^L^ | **0.53** | **<0.001** |
| Chlorophyll content | 0.18 | 0.257 |
| **EWT** ^L^ | **0.51** | **<0.001** |
| **δ13C** | **-0.38** | **<0.05** |
| δ15N | -0.06 | 0.666 |
| Leaf N | 0.19 | 0.257 |
